# Disease-specific fibroblast-myeloid interactions in rheumatoid arthritis synovium

**DOI:** 10.1101/2025.11.17.688477

**Authors:** Alexandra Khmelevskaya, Katerina Apostolopoulou, Camino Calvo Cebrian, Ege Ezen, Melpomeni Toitou, Phatthamon Laphanuwat, Masoumeh Mirrahimi, Celina Geiss, Marija Lugar, Asimina Kakale, Silja Malkewitz, Andrea Laimbacher, Dimitris Konstantopoulos, Vagelis Rinotas, Marietta Armaka, Miranda Houtman, Eva Meier, Kristina Bürki, Justine Leclerc, Hashem Mohammadian, Christof Seiler, Eva Camarillo-Retamosa, Chantal Pauli, Andreas Ramming, Ursula Fearon, Katharina Zachariassen, Thomas Rauer, Adrian Ciurea, Muriel Elhai, Raphael Micheroli, Caroline Ospelt

## Abstract

Rheumatoid arthritis (RA) is characterized by profound remodeling of the synovial microenvironment. Here we show that enhanced fibroblast–macrophage cross-talk distinguishes RA from psoriatic arthritis (PsA). MerTK⁻SPP1⁺ macrophages represent the dominant inflammatory myeloid population in RA, interacting with expanded fibroblast subsets through SPP1-mediated signaling. Lining fibroblasts display induction of antigen-presentation and IL-6/JAK–STAT pathways, while a CHI3L1-producing fibroblast population arises specifically in RA and may act as a source of autoantigens. These stromal populations interact closely with FABP5⁺ iDC3 cells and T cells within a disrupted synovial lining, creating a niche driving adaptive immune activation. In contrast, PsA exhibits increased fibroblast– endothelial interactions without major endothelial transcriptional changes. Our data identify SPP1 signaling and fibroblast–myeloid–dendritic interactions as core drivers of RA synovial inflammation that links innate immune activation to the initiation of autoimmunity.

**Graphical abstract:** 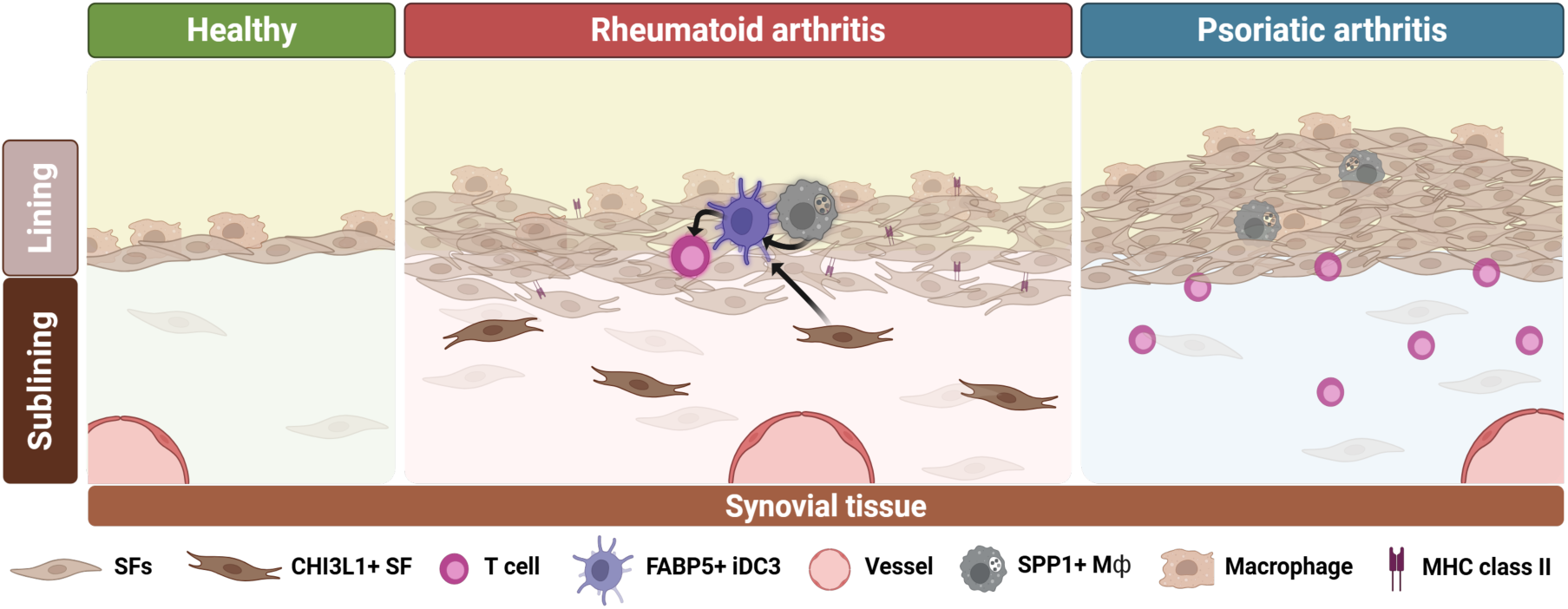

## Introduction

Rheumatoid Arthritis (RA) is a chronic, destructive joint disease that affects around 1% of the world’s population. Local stromal cells of the joint synovial tissue, i.e. synovial fibroblasts, form a complex network with incoming immune cells in RA. Recent in-depth analyses of synovial tissue biopsies have provided groundbreaking insights into the signaling pathways and cells activated in chronic synovitis. Peripheral T helper cells have been identified in the synovial tissue of RA as key cells promoting the local immune response within the joint. Different dendritic cell (DC) populations could be characterized as tolerogenic in healthy synovium, whereas in RA they activate effector memory T cells and naïve T cells^1^. In addition, different macrophage subtypes have been discovered in synovial tissue, with MerTK^-^ macrophages exerting proinflammatory effects, while MerTK^+^ macrophages contribute to the resolution of inflammation and maintenance of remission^2^. In addition, several subtypes of synovial fibroblasts have been identified in RA, which can be divided into invasively growing lining populations and proinflammatory sublining populations. Sublining synovial fibroblast populations are expanded in RA, and single-cell RNA sequencing analyses have suggested multiple functionally distinct cell states^3,4^. The causes of these diverse fibroblast phenotypes are not yet fully understood, but they are thought to be influenced by tissue architecture and interactions with other synovial cell types. For example, endothelial Notch signaling has been shown to stimulate the development of a specific perivascular synovial fibroblast phenotype^5^. Cultured RA synovial fibroblasts lose these different phenotypes but have been shown to be epigenetically imprinted and to still invade and destroy cartilage through the production of matrix metalloproteinases after several passages in culture^6^.

Despite increasing knowledge about the activation of synovial cells in RA, some important questions remain unanswered. It is unclear whether the activation of fibroblasts and immune cell populations is specific to RA or a general feature of chronic synovitis, as most studies have compared RA with degenerative joint diseases (osteoarthritis, OA) or healthy synovia. Furthermore, the functional roles and origins of most fibroblast subpopulations are unknown, as are their interactions with infiltrating immune cell populations. To better understand the cellular landscape and signaling dynamics across chronic arthritides, we compared cellular states and pathways to distinguish a shared inflammatory synovial response from disease-specific mechanisms. Our analysis revealed that macrophage-macrophage interactions, particularly those involving SPP1⁺ macrophages, and macrophage-DC interactions were key features that differentiated RA from psoriatic arthritis (PsA) synovitis. Additionally, fibroblast populations exhibited greater diversification in RA than in other conditions studied. These fibroblasts were characterized by the expression of a previously^7,8^ identified RA autoantigen (CHI3L1) and activation of antigen-presentation pathways.

## Results

### RA synovium is characterised by increased myeloid cell interactions and expansion of synovial fibroblast populations

To characterize the cellular landscape of synovial inflammation across arthritides, we performed ultrasound-guided synovial biopsies from inflamed joints of patients with rheumatoid arthritis (RA; n = 13), psoriatic arthritis (PsA; n = 12), undifferentiated arthritis (UA; n = 5), and osteoarthritis (OA; n = 5). Synovial tissue from individuals with traumatic joint injury without inflammatory arthritis (n = 6) served as non-inflammatory controls (NIC) (Table 1, 2). The cohort was balanced with respect to inflammatory burden, as assessed by the Krenn synovitis score, histopathological classification, and circulating C-reactive protein (CRP) levels. Nonetheless, disease-specific features were evident: PsA patients were predominantly male, RA patients more frequently seropositive, and a greater proportion of RA patients were receiving disease-modifying antirheumatic drugs (DMARDs).

**Table 1.** Clinical characteristics of the patients in scRNAseq cohort. RA = rheumatoid arthritis, PsA = psoriatic arthritis; UA = undifferentiated arthritis; OA = osteoarthritis; NIC = non-inflammatory control; F = female; M = male; seropositivity = positive for rheumatoid factor and/or anti-CCP antibodies; GC = glucocorticoids; MTX = methotrexate; TNFi = TNF inhibitors; JAKi = Janus kinase inhibitors; IL23i = IL-23 inhibitors; CTLA4i = CTLA4 inhibitors; SSZ = sulfasalazine.

|  | Total | RA | PsA | UA | OA | NIC | p-value |
| --- | --- | --- | --- | --- | --- | --- | --- |
|  | (n=41) | (n=13) | (n=12) | (n=5) | (n=5) | (n=6) |  |
| Age (years) |  |  |  |  |  |  |  |
| Mean (SD) | 52 (±16) | 57 (±14) | 49 (±18) | 50 (±3.7) | 74 (±10) | 41 (±16) | 0.033 (ANOVA) |
| Missing | 2 (4.9%) | 0 (0%) | 0 (0%) | 0 (0%) | 2 (40.0%) | 0 (0%) |  |
| Sex |  |  |  |  |  |  |  |
| F | 27 (66 %) | 12 (92 %) | 7 (58 %) | 5 (100 %) | 2 (40 %) | 1 (17 %) | 0.005 (Pearson's Chi-squared test) |
| M | 14 (34 %) | 1 (8 %) | 5 (42 %) | 0 (0 %) | 3 (60 %) | 5 (83 %) |  |
| BMI |  |  |  |  |  |  |  |
| Mean (SD) | 25 (±5.2) | 24 (±5.3) | 26 (±5.0) | NA | NA | NA | 0.226 (ANOVA) |
| Missing | 21 (51.2%) | 4 (30.8%) | 1 (8.3%) | 5 (100%) | 5 (100%) | 6 (100%) |  |
| Seropositivity |  |  |  |  |  |  |  |
| seronegative | 20 (49 %) | 3 (23 %) | 12 (100 %) | 5 (100 %) | NA | NA | 0.0006 (Pearson's Chi-squared test) |
| seropositive | 9 (22 %) | 9 (69 %) | 0 (0 %) | 0 (0 %) | NA | NA |  |
| Missing | 12 (28.8%) | 1 (8 %) | 0 (0%) | 0 (0%) | 5 (100%) | 6 (100%) |  |
| HLAB27 |  |  |  |  |  |  |  |
| neg | 23 (56 %) | 10 (77 %) | 9 (75 %) | 4 (80 %) | NA | NA | 0.591 (Pearson's Chi-squared test) |
| pos | 1 (2 %) | 0 (0 %) | 1 (8 %) | 0 (0 %) | NA | NA |  |
| Missing | 17 (41.0%) | 3 (23.4%) | 2 (16.7%) | 1 (20.0%) | 5 (100%) | 6 (100%) |  |
| Number of affected joints |  |  |  |  |  |  |  |
| Mean (SD) | 7.8 (±5.0) | 8.6 (±4.8) | 8.5 (±5.2) | 4.2 (±4.0) | NA | NA | 0.206 (ANOVA) |
| Missing | 12 (29.3%) | 0 (0%) | 1 (8.3%) | 0 (0%) | 5 (100%) | 6 (100%) |  |
| Number of previous treatments |  |  |  |  |  |  |  |
| Mean (SD) | 3.3 (±3.7) | 4.7 (±3.3) | 3.2 (±4.3) | 0.40 (±0.55) | NA | NA | 0.095 (ANOVA) |
| Missing | 12 (29.3%) | 1 (7.7%) | 0 (0%) | 0 (0%) | 5 (100%) | 6 (100%) |  |
| GC treatment |  |  |  |  |  |  |  |
| no | 21 (51 %) | 9 (69 %) | 9 (75 %) | 3 (60 %) | NA | NA | 0.767 (Pearson's Chi-squared test) |
| yes | 8 (20 %) | 3 (23 %) | 3 (25 %) | 2 (40 %) | NA | NA |  |
| Missing | 12 (28.8%) | 1 (8 %) | 0 (0%) | 0 (0%) | 5 (100%) | 6 (100%) |  |
| DMARD treatment |  |  |  |  |  |  |  |
| MTX | 5 (12 %) | 0 (0 %) | 4 (33 %) | 1 (20 %) | NA | NA | 0.003 (Pearson's Chi-squared test) |
| TNFi | 6 (15 %) | 6 (46 %) | 0 (0 %) | 0 (0 %) | NA | NA |  |
| JAKi | 3 (7 %) | 3 (23 %) | 0 (0 %) | 0 (0 %) | NA | NA |  |
| IL23i | 2 (5 %) | 0 (0 %) | 2 (17 %) | 0 (0 %) | NA | NA |  |
| CTLA4i | 2 (5 %) | 2 (15 %) | 0 (0 %) | 0 (0 %) | NA | NA |  |
| SSZ | 1 (2 %) | 0 (0 %) | 0 (0 %) | 1 (20 %) | NA | NA |  |
| no | 10 (24 %) | 1 (8 %) | 6 (50 %) | 3 (60 %) | NA | NA |  |
| Missing | 12 (28.8%) | 1 (8 %) | 0 (0%) | 0 (0%) | 5 (100%) | 6 (100%) |  |
| Rheumatoid factor |  |  |  |  |  |  |  |
| Mean (SD) | 21 (±62) | 50 (±90) | 0 (±0) | 0 (±0) | NA | NA | 0.095 (ANOVA) |
| Missing | 12 (29.3%) | 1 (7.7%) | 0 (0%) | 0 (0%) | 5 (100%) | 6 (100%) |  |
| anti-CCP antibodies |  |  |  |  |  |  |  |
| Mean (SD) | 50 (±130) | 120 (±180) | 0 (±0) | 0 (±0) | NA | NA | 0.032 (ANOVA) |
| Missing | 12 (29.3%) | 1 (7.7%) | 0 (0%) | 0 (0%) | 5 (100%) | 6 (100%) |  |
| CRP |  |  |  |  |  |  |  |
| Mean (SD) | 15 (±17) | 20 (±20) | 11 (±15) | 10 (±17) | 0.9 | NA | 0.456 (ANOVA) |
| Missing | 10 (24.4%) | 0 (0%) | 0 (0%) | 0 (0%) | 4 (80.0%) | 6 (100%) |  |

**Table 1.1.** Clinical characteristics of the patients in spatial transcriptomics cohort. RA = rheumatoid arthritis, PsA = psoriatic arthritis; OA = osteoarthritis; F = female; M = male; GC = glucocorticoids; HCQ = hydroxychloroquine, SSZ = sulfasalazine; seropositivity = positive for rheumatoid factor and/or anti-CCP antibodies.

|  | Total | RA | PsA | OA |
| --- | --- | --- | --- | --- |
|  | (n=4) | (n=2) | (n=1) | (n=1) |
| Age (years) |  |  |  |  |
| Mean (SD) | 60 (±13) | 66 (±1.4) | 41 | 68 |
| Sex |  |  |  |  |
| F | 2 (50%) | 1 (50 %) | 0 (0%) | 1 (100 %) |
| M | 2 (50%) | 1 (50 %) | 1 (100%) | 0 (0%) |
| GC treatment |  |  |  |  |
| no | 1 (25%) | 0 (0%) | 1 (100%) | NA |
| yes | 2 (50%) | 2 (100%) | 0 (0%) | NA |
| Missing | 1 (25%) | 0 (0%) | 0 (0%) | 1 (100%) |
| DMARD treatment |  |  |  |  |
| HCQ | 2 (50%) | 2 (100%) | 0 (0%) | NA |
| SSZ | 1 (25%) | 1 (50 %) | 0 (0%) | NA |
| no | 1 (25%) | 0 (0%) | 1 (100%) | NA |
| Missing | 1 (25%) | 0 (0%) | 0 (0%) | 1 (100%) |
| Seropositivity |  |  |  |  |
| Positive | 1 (25%) | 1 (50%) | 0 (0%) | NA |
| Negative | 1 (25%) | 0 (0%) | 1 (100%) | NA |
| Missing | 2 (50%) | 1 (50%) | 0 (0%) | 1 (100%) |

**Table 2.** Histological characteristics of the synovial biopsies in scRNAseq cohort. RA = rheumatoid arthritis, PsA = psoriatic arthritis; UA = undifferentiated arthritis; OA = osteoarthritis; NIC = non-inflammatory control; MCP = metacarpophalangeal joint; MTP = metatarsophalangeal joint; PIP = proximal interphalangeal joint.

|  | Total | RA | PsA | UA | OA | NIC | p-value |
| --- | --- | --- | --- | --- | --- | --- | --- |
|  | (n=43) | (n=15) | (n=12) | (n=5) | (n=5) | (n=6) |  |
| Joint |  |  |  |  |  |  |  |
| Knee | 24 (56 %) | 5 (33 %) | 6 (50 %) | 4 (80 %) | 3 (60 %) | 6 (100 %) | 0.258 (Pearson's<br>Chi-squared test) |
| MCP | 7 (16 %) | 3 (20 %) | 3 (25 %) | 0 (0 %) | 1 (20 %) | 0 (0 %) |  |
| MTP | 1 (2 %) | 0 (0 %) | 1 (8 %) | 0 (0 %) | 0 (0 %) | 0 (0 %) |  |
| PIP | 1 (2 %) | 0 (0 %) | 0 (0 %) | 0 (0 %) | 1 (20 %) | 0 (0 %) |  |
| Wrist | 9 (21 %) | 6 (40 %) | 2 (17 %) | 1 (20 %) | 0 (0 %) | 0 (0 %) |  |
| Missing | 1 (2 %) | 1 (7 %) | 0 (0 %) | 0 (0 %) | 0 (0 %) | 0 (0 %) |  |
| Pathotype |  |  |  |  |  |  |  |
| diffuse-myeloid | 7 (16 %) | 3 (20 %) | 2 (17 %) | 2 (40 %) | NA | NA | 0.836 (Pearson's<br>Chi-squared test) |
| fibroid | 10 (23 %) | 4 (27 %) | 4 (33 %) | 2 (40 %) | NA | NA |  |
| lympho-myeloid | 12 (28 %) | 7 (47 %) | 4 (33 %) | 1 (20 %) | NA | NA |  |
| Missing | 14 (32.2%) | 1 (7 %) | 2 (16.7%) | 0 (0%) | 5 (100%) | 6 (100%) |  |
| Histological synovitis score |  |  |  |  |  |  |  |
| Mean (SD) | 4.1 (±1.7) | 4.7 (±1.3) | 3.5 (±1.6) | 4.0 (±2.3) | NA | NA | 0.252 (ANOVA) |
| Missing | 17 (39.5%) | 4 (26.7%) | 1 (8.3%) | 1 (20.0%) | 5 (100%) | 6 (100%) |  |
| CD15+ cells |  |  |  |  |  |  |  |
| no | 14 (33 %) | 3 (20 %) | 9 (75 %) | 2 (40 %) | NA | NA | 0.087 (Pearson's<br>Chi-squared test) |
| yes | 16 (37 %) | 10 (67 %) | 3 (25 %) | 3 (60 %) | NA | NA |  |
| Missing | 13 (29.9%) | 2 (13.7%) | 0 (0%) | 0 (0%) | 5 (100%) | 6 (100%) |  |
| Vascularisation |  |  |  |  |  |  |  |
| <10 | 15 (35 %) | 7 (47 %) | 8 (67 %) | 0 (0 %) | NA | NA | 0.096 (Pearson's<br>Chi-squared test) |
| >10 | 15 (35 %) | 6 (40 %) | 4 (33 %) | 5 (100 %) | NA | NA |  |
| Missing | 13 (29.9%) | 2 (13.7%) | 0 (0%) | 0 (0%) | 5 (100%) | 6 (100%) |  |
| Lymphoid follicle |  |  |  |  |  |  |  |
| no | 22 (51 %) | 7 (47 %) | 10 (83 %) | 5 (100 %) | NA | NA | 0.185 (Pearson's<br>Chi-squared test) |
| yes | 8 (19 %) | 6 (40 %) | 2 (17 %) | 0 (0 %) | NA | NA |  |
| Missing | 13 (29.9%) | 2 (13.7%) | 0 (0%) | 0 (0%) | 5 (100%) | 6 (100%) |  |
| Fibrosis level |  |  |  |  |  |  |  |
| high | 1 (2 %) | 0 (0 %) | 0 (0 %) | 1 (20 %) | NA | NA | 0.2149<br>(Pearson's Chi-squared test) |
| intermediate | 2 (5 %) | 2 (13 %) | 0 (0 %) | 0 (0 %) | NA | NA |  |
| low | 20 (47 %) | 10 (67 %) | 8 (67 %) | 2 (40 %) | NA | NA |  |
| no | 2 (5 %) | 1 (7 %) | 1 (8 %) | 0 (0 %) | NA | NA |  |
| Missing | 18 (42.2%) | 2 (13.7%) | 3 (25.3%) | 2 (40 %) | 5 (100%) | 6 (100%) |  |

**Table 2.1.** Histological characteristics of the synovial biopsies in spatial transcriptomics cohort. RA = rheumatoid arthritis, PsA = psoriatic arthritis; OA = osteoarthritis.

|  | Total | RA | PsA | OA |
| --- | --- | --- | --- | --- |
|  | (n=4) | (n=2) | (n=1) | (n=1) |
| Joint |  |  |  |  |
| Knee | 3 (75%) | 1 (50 %) | 1 (100%) | 1 (100%) |
| Hip | 1 (25%) | 1 (50 %) | 0 (0%) | 0 (0%) |
| Pathotype |  |  |  |  |
| diffuse-myeloid | 0 (0%) | 0 (0%) | 0 (0%) | 0 (0%) |
| fibroid | 0 (0%) | 0 (0%) | 0 (0%) | 0 (0%) |
| lympho-myeloid | 2 (50%) | 1 (50 %) | 1 (100%) | 0 (0%) |
| Missing | 2 (50%) | 1 (50 %) | 0 (0%) | 1 (100%) |
| Histological synovitis score |  |  |  |  |
| Mean (SD) | 6.5 (±2.1) | 8 | 5 | NA |
| Missing | 2 (50%) | 1 (50 %) | 0 (0%) | 1 (100%) |

We identified the following synovial cell populations based on the expression of canonical marker genes in single-cell RNA sequencing (scRNA-seq) of synovial biopsies (n = 162’833 cells; mean = 3’786 cells per sample) (Fig. 1a, Extended Data Fig. 1a): synovial fibroblasts (*COL1A1*, *THY1*, *PRG4*, etc.), monocyte/macrophages (*LYZ*, *C1QA*, *FCGR3A*, etc.), T/NK cells (*CD3D*, *CD3E*, etc.), endothelial cells (*VWF*, *CCL14*, *ACKR1*, etc.), neutrophils (*S100A8*, *S100A9*, *FCGR3B*, etc.), smooth muscle cells (*ACTA2*, *MYL9*, etc.), dendritic cells (DCs) (*CLEC10A*, *FCER1A*, etc.), B cells (*CD79A*, *MS4A1*, *IGKC*, etc.), proliferating cells (*STMN1*, *MKI67*, etc), plasma cells (*MZB1*, *JCHAIN*, etc.), mast cells (*TPSB2*, *TPSAB1*, *CPA3*, etc), and lymphatic endothelial cells (*CCL21*, *RELN*, *TFF3*, etc.). As expected, inflammatory arthritis samples (RA, PsA, UA) exhibited increased immune cell infiltration— particularly T cells, B cells, and neutrophils—relative to OA and NIC tissues (Fig. 1b, Extended Data Fig. 1b,c). Fibroblast abundance was increased in PsA and OA compared to RA, whereas in RA synovium dendritic cells were enriched (Fig. 1b, Extended Data Fig. 1b,d). Fibroblasts and myeloid cells also showed the highest number of genes differentially expressed between RA and PsA (Extended Data Fig. 1e).

**Figure 1.**
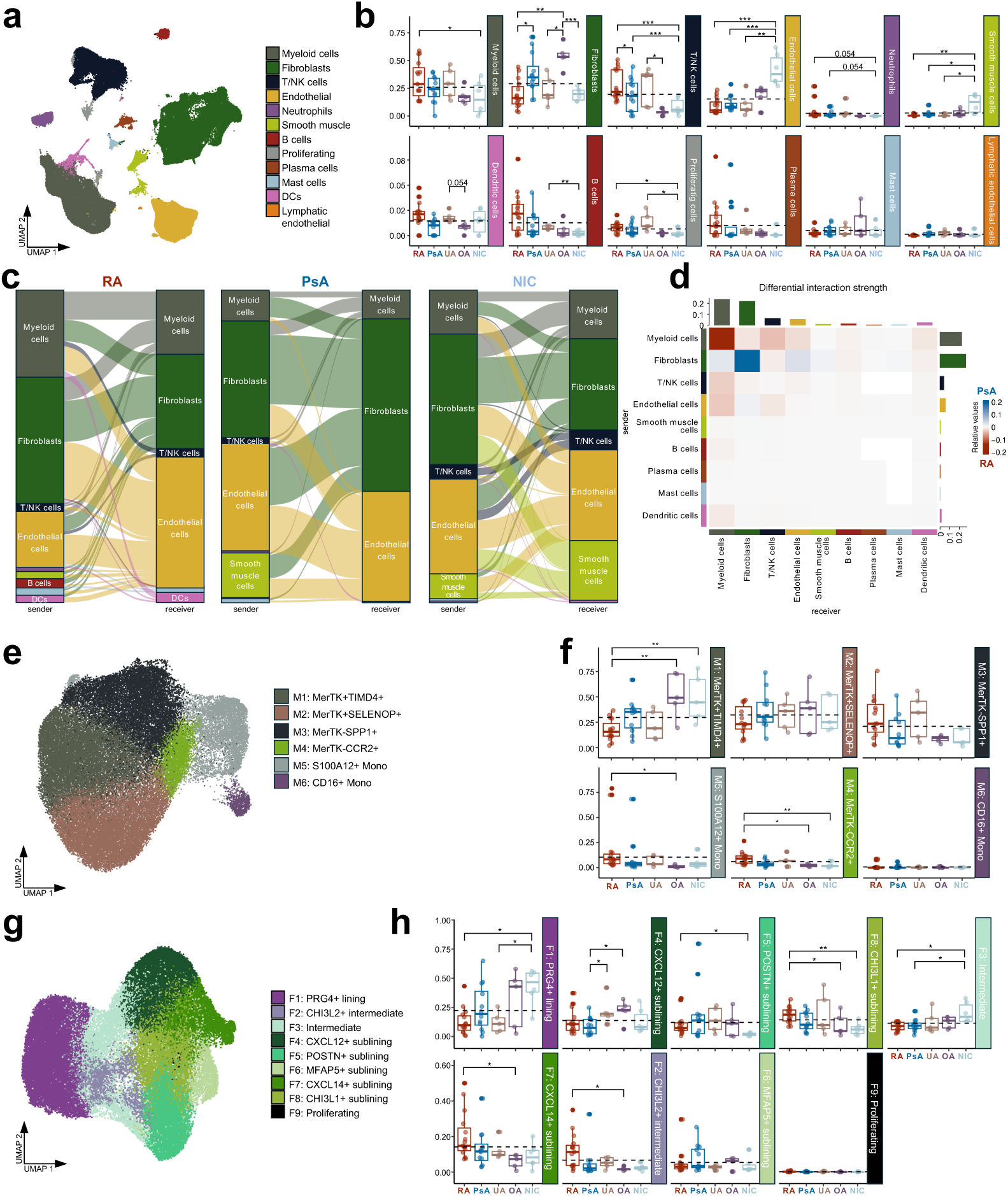
**a** UMAP of main synovial cell populations in unintegrated merged scRNAseq data. **b** Boxplot of the proportions of the main cell populations between diagnoses. n = 15 RA, n = 12 PsA, n = 5 UA, n = 5 OA, n = 6 NIC. **c** Alluvial plot of top 250 interactions in RA, PsA, and healthy synovium according to MultinichnetR^32^. **d** Heatmap of differential interaction strength in PsA versus RA according to CellChat^33^. **e** UMAP of myeloid cell populations in integrated scRNAseq data. **f** Boxplot of the proportions of the myeloid cell populations between diagnoses. n = 15 RA, n = 11 PsA, n = 5 UA, n = 5 OA, n = 5 NIC. **g** UMAP of synovial fibroblast populations in integrated scRNAseq data. **h** Boxplot of the proportions of the synovial fibroblast populations between diagnoses. n = 15 RA, n = 12 PsA, n = 5 UA, n = 5 OA, n = 6 NIC. *For each boxplot: Dashed line represents the mean percentage of the populations across diagnoses. The lower and upper hinges correspond to the first and third quartiles (the 25^th^ and 75^th^ percentiles). The upper whisker extends from the hinge to the largest value no further than 1.5 * IQR from the hinge (where IQR is the inter-quartile range, or distance between the first and third quartiles). The lower whisker extends from the hinge to the smallest value at most 1.5 * IQR of the hinge. Differential composition analysis (DCATS^34^) was used with Benjamini & Hochberg p-value adjustment.

Cell-cell communication analysis showed a pronounced increase in myeloid cell interactions in RA, both in terms of frequency (Fig. 1c) and interaction strength, relative to PsA, OA, and NIC tissues (Fig. 1d, Extended Data Fig. 1f,g). In contrast, fibroblast-mediated interactions predominated in PsA and OA synovium (Fig. 1c,d, Extended Data Fig. 1f). Spatial transcriptomic profiling (Extended Data Fig. 1h-j) corroborated these findings, with increased interaction strength observed in SPP1⁺ macrophage cluster in RA compared to PsA (Extended Data Fig. 1k).

To further dissect this RA specific increase in myeloid cell interactions, we subclustered synovial myeloid cells in our scRNAseq data (total 47’178 cells, mean 1’151 myeloid cells per synovial biopsy) after integration with myeloid cells from PBMCs of RA (n = 4) and PsA patients (n = 5) (total 3’967 cells) (Extended Data Fig. 2a,b). Clustering identified two subpopulations of MerTK^+^ macrophages (TIMD4^+^ and SELENOP^+^), two subpopulations of MerTK^-^ macrophages (SPP1^+^ and CCR2^+^) and two monocyte subpopulations (S100A12^+^ and CD16^+^) (Fig. 1e, Extended Fig. 2c,d). As expected, synovial tissue was enriched for macrophages, whereas PBMCs contained a higher proportion of monocytes (Extended Data Fig. 2b,c). Inflamed synovium had an increased influx of monocyte-derived MerTK^-^ SPP1^+^ and MerTK^-^CCR2^+^ macrophages and S100A12^+^ monocytes (Extended Data Fig. 2e,f), which was more pronounced in RA compared to PsA synovial tissues (Fig. 1f).

We also sub-clustered synovial fibroblasts (total of 52’392 cells, mean of 1’218 fibroblasts per synovial biopsy) using unsupervised clustering into a lining fibroblasts subpopulation (PRG4^+)^, two intermediate subpopulations (CHI3L2^+^ and intermediate), five subpopulations of sublining fibroblasts (CXCL12^+^, POSTN^+^, MFAP5^+^, CXCL14^+^ and CHI3L1^+^ sublining) and a subpopulation of proliferating fibroblasts (Fig. 1g, Extended Data Fig. 2g-i). Comparative analysis of fibroblast subset frequencies revealed a significant enrichment of CHI3L1⁺, CXCL14⁺, POSTN⁺, and CHI3L2⁺ fibroblasts in RA relative to OA and NIC tissues (Fig. 1h, Extended Data Fig. 2j). Moreover, CHI3L1⁺, CXCL14⁺, and CHI3L2⁺ subsets were also significantly expanded in RA compared to PsA, whereas POSTN⁺ fibroblasts did not differ significantly between these two conditions (Fig. 1h, Extended Data Fig. 2k).

Together, these data indicated enhanced myeloid cell communication in RA compared to PsA synovium and selective expansion of fibroblast subpopulations supporting the notion of a distinct stromal remodeling program in RA synovium.

### SPP1 is highly expressed in RA monocyte-derived macrophages and induces pro-inflammatory and metabolic changes in macrophages

Despite comparable overall macrophage frequencies across UA, RA, and PsA synovial tissues, transcriptomic analysis revealed that *SPP1* (encoding secreted phosphoprotein 1, also known as osteopontin, OPN) was among the most significantly upregulated genes in RA compared to PsA synovium, with high log fold change and statistical significance (Fig. 2a). *SPP1* expression was confined to MerTK⁻SPP1⁺ macrophages (Extended Data Fig. 3a,b). Plasma concentrations of SPP1 were higher in RA patients compared to axial spondyloarthritis (axSpA), but not to PsA patients (Fig. 2b). On the other hand, there was a statistical trend toward increased levels of SPP1 in RA synovial fluid compared to PsA (Fig. 2c), supporting the notion of RA-specific production of SPP1 predominantly locally in the joints. Signalling via SPP1 was the most differential cell-cell interaction pathway across all cell types in RA versus PsA (Fig. 2d). Notably, hypoxia-related pathways were the most up-regulated in pathway analysis^9^ comparing RA MerTK⁻SPP1⁺ macrophages to PsA (Fig. 2e-f) and non-inflamed synovium (Extended Data Fig. 3c, e-f). Pathway analysis of predicted SPP1 target genes up-regulated in RA MerTK⁻SPP1⁺ macrophages relative to PsA indicated hypoxia-like gene expression induction by SPP1 (Fig. 2g, Extended Data Fig. 3d). Given the established link between hypoxia and glycolytic reprogramming^10^, we hypothesized that elevated SPP1 levels in RA may drive metabolic shifts in macrophages. Indeed, in vitro stimulation of monocyte-derived macrophages (MDMs) with SPP1 resulted in increased lactate production (Fig. 2h), with no change in mitochondrial membrane potential (Fig. 2i) indicating an SPP1-induced switch to aerobic glycolysis. Thus, increased macrophage interactions in RA were mainly driven by SPP1 which in addition to an inflammatory response might mediate metabolic reprogramming of synovial macrophages in RA.

**Figure 2.**
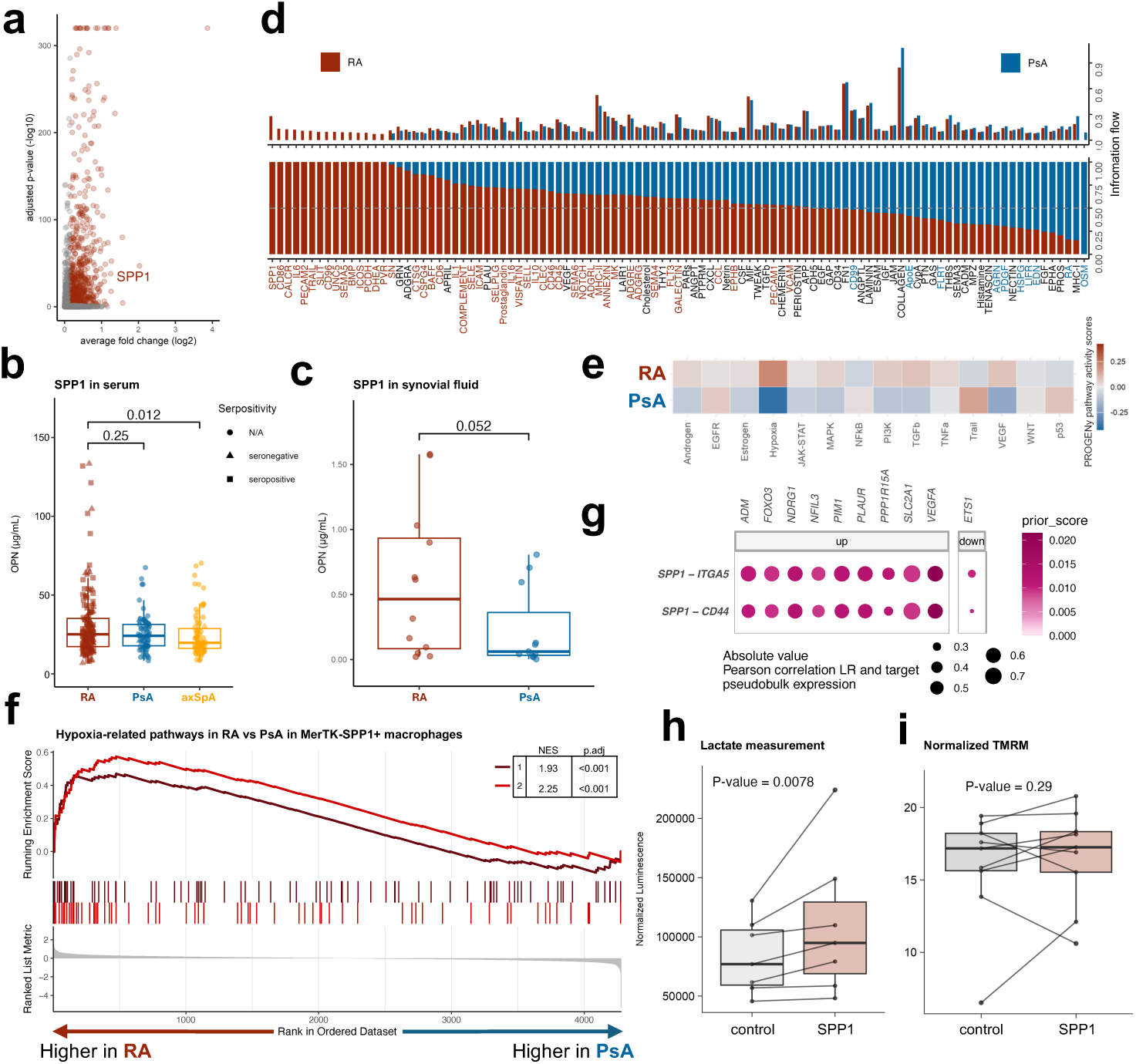
**a** Volcano plot of DEGs in RA versus PsA across all cell types. **b** OPN protein (SPP1) measurements in serum (n = 311: RA = 159, PsA = 72, axSpA = 80) (one-tailed Wilcoxon test). **c** OPN protein (SPP1) measurements in synovial fluid (n = 20: RA = 10, PsA = 10) (right panel) (one-tailed Wilcoxon test). **d** Barplot of information flow of signaling pathways between all cell types in RA versus PsA. **e** PROGENy^9^ pathway activity scores in MerTK-SPP1+ macrophages in RA versus PsA. **f** GSEA results for pathways up-regulated in MerTK-SPP1+ macrophages in RA versus PsA (1 – Hallmark: Hypoxia, 2 – GO: Cellular response to oxygen levels). **g** Prioritization score and Pearson correlation between target genes from “Hallmark: Hypoxia” pathway predicted to be up- or down-regulated in MerTK-SPP1+ macrophages upon SPP1 interaction with ITGA5 and CD44. **h** Normalized luminescence values reflecting lactate concentration in MDM supernatants after SPP1 stimulation (n = 7) and control (n = 7) (one-tailed paired Wilcoxon test). **i** Normalized TMRM fluorescence measured by CX7 in MDM after SPP1 stimulation (n = 9) and control (n = 9) (one-tailed paired T-test).

### RA synovium harbours activated DC populations that interact with macrophages and receive signals from stromal cells

To further dissect the role of MerTK⁻SPP1⁺ macrophages in RA synovial inflammation, we analyzed their cell-cell communication patterns. MerTK⁻SPP1⁺ macrophages predominantly signaled to other macrophages, particularly within the same MerTK⁻SPP1⁺ subset (Fig. 3a). However, this predominant signal of MerTK⁻SPP1⁺ macrophages to other macrophages was not disease specific. In contrast, signals sent from MerTK⁻SPP1⁺ macrophage to DCs (Fig. 3a) as well as signals received by MerTK⁻SPP1⁺ macrophages from SFs (specifically CHI3L1^+^, CXCL12 ^+^ and CXCL14 ^+^ sublining fibroblasts) were increased in RA synovium (Fig. 3b). Further analysis of DC subsets identified eight DC subpopulations, including FABP5^+^ inflammatory DC3 (FABP5^+^ iDC3) (Fig. 3c, Extended Data Figure 4a-d). This iDC3 population was previously shown to have a crucial role in promoting T cell activation in RA synovium^1^. While the relative abundance of DC subsets was comparable across disease groups (Fig. 3c, Extended Data Fig. 4e), differential gene expression analysis showed that FABP5⁺ iDC3s exhibited the highest number of RA-specific transcriptional changes compared to PsA (Fig. 3d). Moreover, this subset demonstrated the greatest number of predicted cell-cell interactions in RA, but not in PsA (Fig. 3e), with incoming signals primarily originating from MerTK⁻SPP1⁺ macrophages, as well as CHI3L1⁺ fibroblasts (Fig. 3f) and outgoing signals mainly to MerTK⁻SPP1⁺ macrophages (Extended Data Fig. 4h). Analysis of signaling modalities indicated that MerTK⁻SPP1⁺ macrophages interacted with FABP5⁺ iDC3s predominantly through direct cell-cell contact, while CHI3L1^+^ fibroblasts relied mostly on secreted and extracellular matrix–mediated signaling (Fig. 3g). Accordingly, iDC3 had the highest expression of CHI3L1 receptors among DC subpopulations (Extended Data Fig. 4f) and the expression was higher in RA patients in comparison to all other diagnosis (Extended Data Fig. 4g). Spatial transcriptomic data further supported a direct interaction between MerTK⁻SPP1⁺ macrophages and FABP5⁺ iDC3s, showing that regions enriched for FABP5⁺ iDC3s exhibited the highest levels of SPP1 expression, which declined with increasing distance from these regions (Fig. 3h-j). Notably, this spatial association was specific to RA and absent in PsA and OA tissues (Fig. 3j). On the other hand, CHI3L1^+^ fibroblasts did not show a correlation with the distance to iDC3 in any diagnosis (Extended Data Fig. 4i). Multiplex immunofluorescent (IF) staining confirmed the findings from single-cell and spatial transcriptomics analysis, showing direct interactions between SPP1^+^ macrophages and DCs (Fig.3k), as well as with T cells (Extended Data Fig. 4j). At the same time, CHI3L1^+^ fibroblasts were located farther away from DCs, which aligns with transcriptomic analyses demonstrating that fibroblast – DC interactions primarily depend on secreted signaling pathways (Extended Data Fig. 4j).

**Figure 3.**
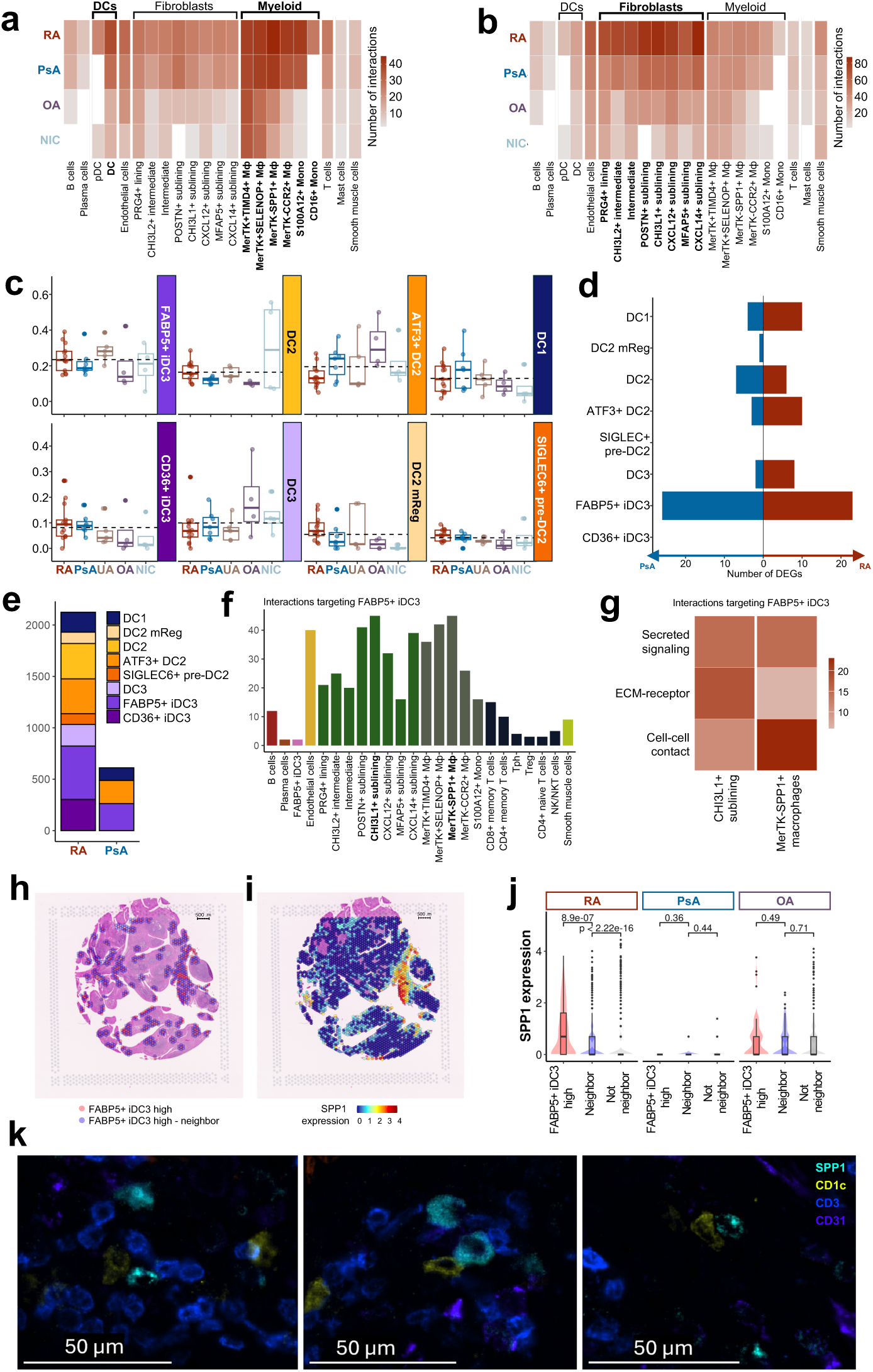
**a** Heatmap of the number of interactions outgoing from MerTK-SPP1+ macrophages with a probability > 8% according to CellChat^33^. **b** Heatmap of the number of interaction incoming to MerTK-SPP1+ macrophages with a probability > 8% according to CellChat^33^. **c** Boxplot of the proportions of DC populations between diagnoses. n = 13 RA, n = 8 PsA, n = 5 UA, n = 4 OA, n = 4 NIC. The dashed line represents mean percentage of the populations across diagnoses. The lower and upper hinges correspond to the first and third quartiles (the 25^th^ and 75^th^ percentiles). The upper whisker extends from the hinge to the largest value no further than 1.5 * IQR from the hinge (where IQR is the inter-quartile range, or distance between the first and third quartiles). The lower whisker extends from the hinge to the smallest value at most 1.5 * IQR of the hinge. **d** Numbers of DEGs (log2FC > 0.15 and p-value adjusted < 0.05) between RA and PsA by DC subpopulation. **e** Number of interactions targeting different dendritic cell populations with probability > 8% according to CellChat^33^ in RA and PsA. **f** Number of interaction targeting FABP5^+^ iDC3 with the probability > 8% in RA according to CellChat^33^. **g** Number of the interactions incoming to FABP5^+^ iDC3 with the probability > 8% in RA by type of signaling according to CellChat^33^. **h** Representative spatial transcriptomics RA slide with annotation of FABP5^+^ iDC3 high spots and their neighbours. **i** Representative spatial transcriptomics RA slide with SPP1 expression values in FABP5^+^ iDC3 high neighbourhoods only. **j** Boxplot with violin plots of SPP1 expression values across FABP5+ iDC3 “neighbourhoods” across diagnosis in spatial transcriptomics data (Wilcox test). **k** Multiplex IF staining of DCs (CD1c-positive), T cells (CD3-positive), SPP1^+^ macrophages (SPP1-positive) and endothelial cells (CD31-positive) in RA synovial tissue.

Together, these findings demonstrated that MerTK⁻SPP1⁺ macrophages promote disease-specific macrophage–DC interactions in RA synovial tissues. Given the expansion of distinct fibroblast subsets in RA, especially of CHI3L1^+^ sublining fibroblasts (Extended Data Fig. 2k), we next sought to determine whether these stromal populations contribute to the activation of myeloid cells in the RA synovial microenvironment.

### CHI3L1 expression is specifically increased and epigenetically imprinted in RA synovial fibroblasts

The appearance of a CHI3L1^+^ fibroblast population in RA and its strong signaling outgoing to MerTK^-^ SPP1^+^ macrophages (Fig. 3b) and to FABP5^+^ iDC3 (Fig. 3f) in RA raised our interest. CHI3L1, also known as YKL-40, chondrex or HC-gp39, is a secreted glycoprotein that has previously been described as autoantigen in RA and to be arthritogenic in mice^8^. Spatial transcriptomic analysis (Fig. 4a) and protein staining (Fig. 4b) localized CHI3L1 to sublining fibroblasts residing just beneath the lining layer. Analysis of differentially expressed genes (DEG) in synovial fibroblasts between diagnoses showed significantly higher expression of *CHI3L1* in RA comparing to PsA (log fold change = 1.66, adj. p-value < 0.001) (Fig. 4c) and non-inflamed tissues (Extended Data Fig. 5a,b). We confirmed the significantly higher expression of *CHI3L1* in RA versus PsA in publicly available data^11^ (Extended Data Fig. 5c) and in mouse models of RA (hTNF-Tg) in comparison to PsA (A20^Znf7-KI^) (Extended Data Fig. 5d-e). Concentrations of CHI3L1 were significantly higher in the serum of RA patients compared to PsA independent of presence of autoantibodies (rheumatoid factor and/or anti-CCP antibodies) (Fig. 4d) and discrimination between RA and PsA diagnosis using logistic regression model based only on CHI3L1 serum levels showed a mean AUC of 0.7123 (Fig. 4e). There was a significant positive correlation between CHI3L1 concentration and age (Extended Data Fig. 5f). Nevertheless, when applying a linear regression model adjusting for sex, age, and time since diagnosis, CHI3L1 concentration remained significantly different between diagnostic groups, with higher levels observed in RA. Furthermore, analysis of variance confirmed that inclusion of diagnosis significantly increased the explanatory power of the model (F = 4.58, p = 0.033), supporting diagnosis as an independent determinant of CHI3L1 concentration. *CHI3L1* expression was increased in several synovial fibroblast populations in RA (Extended Fig. 5g,h) but was absent from any other synovial or blood cell type (Fig. 4f), which makes synovial fibroblasts a likely source for circulating CHI3L1. However, chondrocytes and bone cells were also shown to express CHI3L1 and might contribute to the observed differences in serum levels^12^.

**Figure 4.**
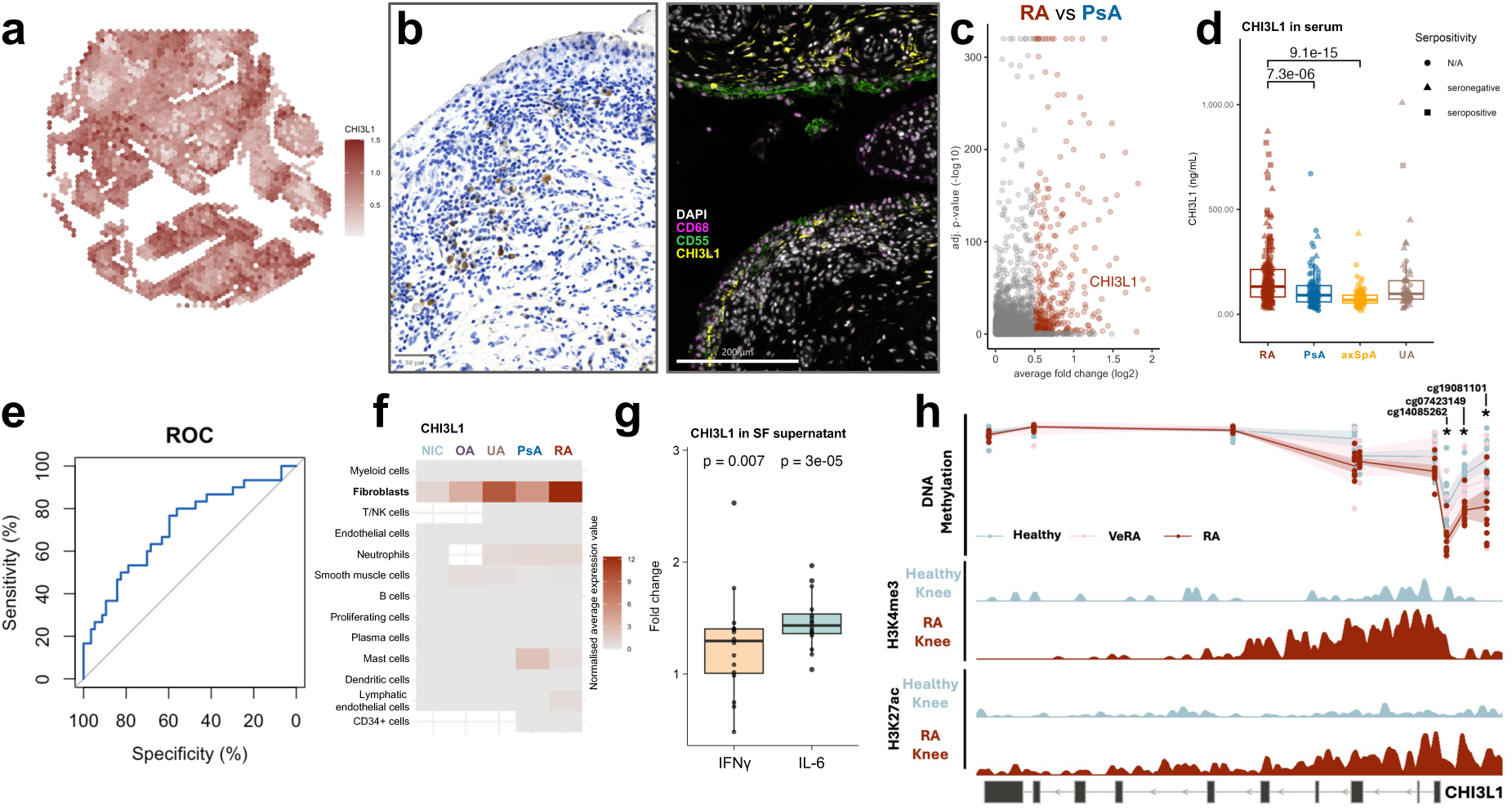
**a** *CHI3L1* gene expression level in RA synovial tissue. **b** Left panel: Immunohistochemical staining of CHI3L1 in RA synovial tissue. Right panel: Multiplex immunofluorescent staining of RA synovial tissue. **c** Volcano plot of DEGs up-regulated in RA versus PsA fibroblasts. **d** Boxplot of CHI3L1 concentration in serum measured by ELISA (Seropositive patients – positive for rheumatoid factor and/or anti-CCP antibodies) (one-tailed Wilcoxon test) (n = 440: RA = 190, PsA = 99, axSpA = 100, UA = 51). **e** ROC curve analysis of the power of CHI3L1 serum levels to discriminate between RA and PsA diagnosis. **f** Heatmap of mean *CHI3L1* expression levels across cell types in synovium and PBMC. **g** Fold change values of CHI3L1 concentration in synovial fibroblast supernatant after stimulation versus control (n = 16) (one-tailed paired Wilcoxon test). **h** Genome tracks with DNA methylation levels (beta values) in healthy (n = 15), very early RA (VeRA, n = 6), and RA (n = 11) samples, along with ChIP-seq coverage for H3K4me3 (marking promoter activity) and H3K27ac (marking active enhancers) from a healthy and an RA donor. Three CpG sites located in the core promoter (cg14085262, cg07423149, and cg19081101) with significantly changed DNA methylation levels are highlighted.

In vitro stimulation with IL-1β, IL-6 and IFNγ, but not TNFα induced secretion of CHI3L1 by cultured synovial fibroblasts (Fig. 4g, Extended Data Figure 5i) irrespective of diagnosis (Extended Data Figure 5j). Consistently, re-analysis of publicly available bulk RNA-seq data of synovial fibroblasts exposed to different stimuli^13^ demonstrated that IL-1β, IFNγ, and IL-6 stimulation led to up-regulation of *CHI3L1* gene expression, whereas TGF-β and TNFα stimulation resulted in its down-regulation (Extended Data Fig. 5k). Finally, supernatants from T cells obtained from RA patients, but not from healthy individuals, significantly induced CHI3L1 production by fibroblasts (Extended Data Fig. 5l)

The CHI3L1 gene locus was previously reported to be hypomethylated in RA versus OA synovial fibroblasts^14^. In our dataset, we observed hypomethylation of CHI3L1 in RA synovial fibroblasts compared to healthy controls. Furthermore, when including synovial fibroblasts from patients with very early RA (VeRA), we observed a gradient of methylation from healthy to VeRA to established RA, suggesting progressive epigenetic remodeling during disease development (Fig. 4h, Extended Data Fig. 5m). Additionally, H3K4me3 promoter mark and H3K27ac active enhancer mark were enriched at the CHI3L1 locus in RA synovial fibroblasts compared to healthy controls. Together, these findings indicate epigenetic imprinting of increased CHI3L1 transcription in RA synovial fibroblasts at very early stages of the disease.

Rather unexpectedly, silencing of CHI3L1 in RA synovial fibroblasts (Extended Data Fig. 6a) led to upregulation of inflammatory response genes (Fig. 5a), particularly those associated with response to IFN (Fig. 5b) with upregulation of *IRF1* after silencing (Extended Data Fig. 6b). On the other hand, extracellular matrix pathways (Fig. 5c) and *COL1A1* (Extended Data Fig. 6c) were downregulated and the expression of the matrix-degrading enzyme *MMP1* increased (Extended Data Fig. 6d). Genes that were increased after silencing of CHI3L1 were also enriched in gene sets measured in synovial fibroblasts after stimulation with IL-6, IL-1, or IFNs (Fig. 5d, Extended Data Fig. 6e). These findings suggest that CHI3L1 is induced as a counter-regulatory mechanism to modulate inflammatory responses. Accordingly, the induction of IFN response genes (*MX2*), inflammatory genes (IL-6) and MMPs was higher in CHI3L1 silenced synovial fibroblasts under stimulation, while collagen production was lower in CHI3L1 silenced fibroblasts compared to control silenced fibroblasts under stimulation (Fig. 5e, Extended Data Fig. 6f-g).

**Figure 5.**
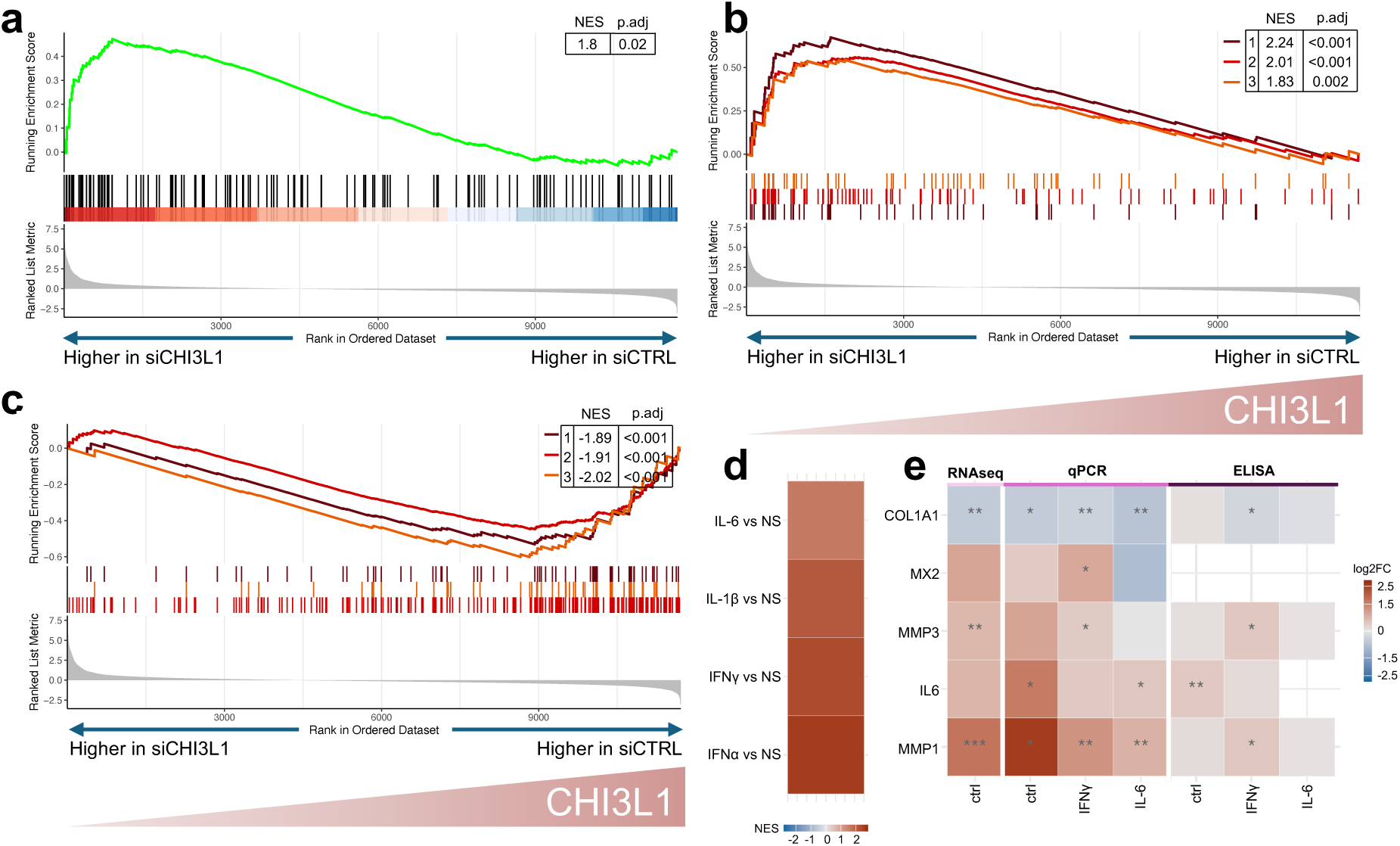
**a** GSEA result for pathway “Hallmark: Inflammatory response” in siCHI3L1 versus siCTRL. **b** GSEA results for pathways up-regulated after silencing (1 – Reactome: IFNα/β signaling, 2 – Hallmark: IFNα response, 3 – GO: Response to type I IFN). **c** GSEA results for pathways down-regulated after silencing (1 – Reactome: Collagen formation, 2 – Reactome: Extracellular matrix organization, 3 – GO: Collagen fibril organization). **d** Heatmap of normalized enrichment scores (NES) of pathways retrieved from Tsuchiya *et al.*^13^ in siCHI3L1 versus siCTRL gene list. **e** Heatmap of log2 fold change values of gene expression and protein concentration in siCHI3L1 versus siCTRL SFs across different types of analysis.

On the other hand, stimulation of synovial fibroblasts with recombinant CHI3L1 did neither influence IFN nor inflammatory response genes under basal nor stimulated conditions (Extended Data Fig. 7a-c), suggesting that counterregulatory CHI3L1 functions are primarily mediated through an endogenous, cell-intrinsic mechanism in synovial fibroblasts, rather than acting via endocrine signaling. Gene set enrichment analysis still showed induction of extracellular matrix organization via CHI3L1 (Extended Data Fig. 7d). Nonetheless, recombinant CHI3L1 did not alter COL1A1 or MMP production under basal conditions (Extended Data Fig. 7c,e,f). Upon co-stimulation with IFNγ, *COL1A1* expression decreased slightly, with an average log2 fold change below 0.25 (Extended Data Fig. 7e,g).

To see which other cells might be affected by increased CHI3L1 levels in the synovium, we performed a predictive interaction analysis of CHI3L1 with its receptors CD44, IL13RA2 and LGALS3 across all synovial cell types. Cell-cell interaction analysis suggested most cell-cell interactions of CHI3L1^+^ sublining fibroblasts with MerTK^-^SPP1^+^ macrophages (Extended Data Fig. 7h). Previously, it was shown that CHI3L1 increased secretion of IL-1β by THP1 monocytic cells^15^. However, we could not confirm this (Extended Data Fig. 7i) and stimulation of MDM with CHI3L1 had no effect on SPP1 production (Extended Data Fig. 7j), as well as on expression of inflammation-associated genes (Extended Data Fig. 7k).

These findings suggest that the induction of CHI3L1 expression in synovial fibroblasts is an early event in the pathogenesis of RA, potentially triggered by an inflammatory response aimed at dampening inflammation. However, the elevated levels of CHI3L1 in the synovium do not appear to directly account for the inflammatory activation of synovial cells. Instead, CHI3L1 may act as an autoreactive factor, potentially contributing to the initiation or amplification of local adaptive immune responses.

### Lining synovial fibroblasts show higher involvement in inflammatory response in RA compared to PsA

Although increased proportions of sublining synovial fibroblasts appeared as a hallmark of RA, the highest number of DEG across all diseases compared to healthy controls was observed in PRG4^+^ lining synovial fibroblasts (Fig. 6a), showing the high level of involvement of these fibroblasts in synovial inflammation irrespective of their decrease in proportional abundance (Fig. 1h, Extended Data Fig. 2j,k). Pathway analysis of genes differentially expressed in PRG4^+^ lining in RA compared to PsA showed that RA lining upregulated inflammatory as well as hypoxia related pathways (Fig. 6b), similar to what we saw in MerTK^-^SPP1^+^ macrophages in RA. Analysis of synovial fibroblasts from a publicly available dataset^11^ confirmed these pathways activated in lining RA fibroblasts compared to PsA (Extended Data Fig. 8a). However, unlike in MDMs (Fig. 4h-i), SPP1 did not change lactate production in cultured synovial fibroblasts (Extended Data Fig. 8b). A metabolic switch in RA lining fibroblasts has previously been reported to be induced by TGF-β and IL-1β in RA fibroblasts^11^. Therefore, changes in cell metabolism in RA appear to be regulated by cell type-specific mechanisms.

**Figure 6.**
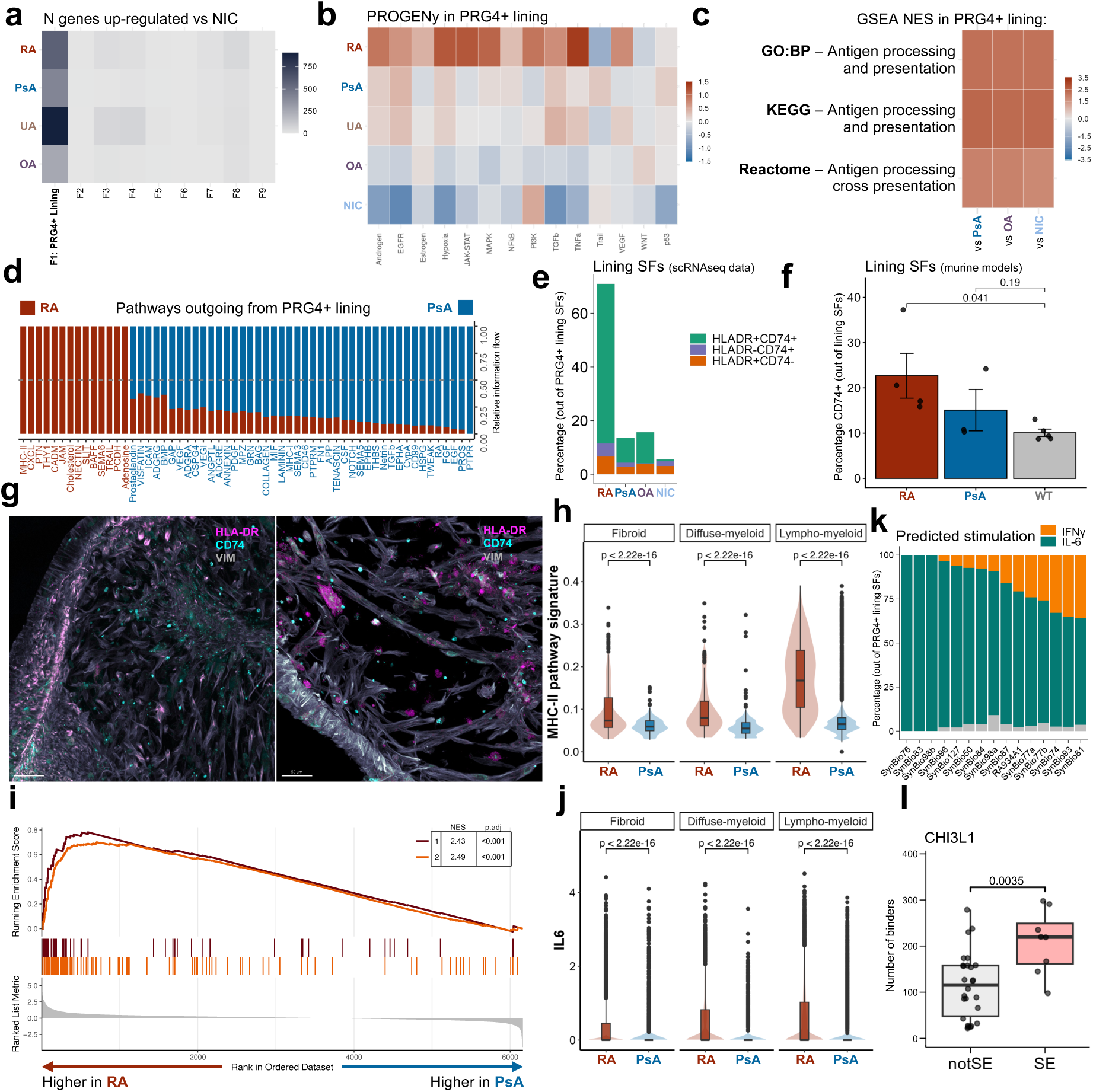
**a** Heatmap of the number of up-regulated DEGs in comparison to healthy controls (NIC). **b** Heatmap of pathway responsive genes for activity inference analysis (PROGENy^9^) in PRG4^+^ lining fibroblasts across diseases. **c** Heatmap of GSEA normalized enrichment score (NES) in PRG4+ lining fibroblasts in RA in comparison to other diagnoses. **d** Barplot of cell-cell interactions outgoing from PRG4^+^ lining fibroblasts in RA compared to PsA. **e** Barplot of the proportion of HLA-DR^+^ and/or CD74^+^ lining synovial fibroblasts across diagnoses. **f** Proportion of CD74+ lining synovial fibroblasts in RA mouse model (hTNF-Tg) (n = 4), PsA mouse model (A20^Znf7-KI^) (n = 3) and healthy mice (wild type, WT) (n = 5) (average ± SEM is shown for each group of mice, * represent p<0.05, one-way ANOVA with Bonferroni correction between indicated genotypes). **g** Confocal multiplex immunofluorescence image of micromasses showing VIM, CD74, and HLA-DR staining. **h** Violin plot of GO:BP term ‘Antigen processing and presentation of exogenous peptide antigen via MHC class II’ signature in fibroid, diffuse-myeloid and lympho-myeloid tissues of RA (n = 4 fibroid, n = 3 diffuse-myeloid, n = 7 lympho-myeloid) and PsA patients (n = 3 fibroid, n = 1 diffuse-myeloid, n = 5 lympho-myeloid) (Wilcox test). **i** GSEA results in PRG4+ lining fibroblasts in RA versus PsA for pathways: 1 - Reactome: Interferon gamma signaling, 2 – Hallmark: Interferon gamma response. **j** Violin plot of *IL6* expression in all synovial fibroblasts in fibroid, diffuse-myeloid and lympho-myeloid tissues of RA (n = 4 fibroid, n = 3 diffuse-myeloid, n = 7 lympho-myeloid) and PsA patients (n = 3 fibroid, n = 1 diffuse-myeloid, n = 5 lympho-myeloid) (Wilcox test). **k** Barplot showing the proportion of cells with predicted stimulations (IL-6, IFNγ, or unclassified (in grey)) within PRG4^+^ lining SFs of RA patients. **l** Boxplot with the numbers of peptides derived from CHI3L1 predicted to bind to HLA class II alleles. SE – shared epitope alleles (Wilcox test).

PRG4⁺ lining fibroblasts had more outgoing interactions in RA in comparison to PsA (Extended Data Fig. 8c) and upregulated genes involved in antigen processing and presentation (Fig. 6c). Cell-cell interaction analysis showed MHC-II pathways significantly up-regulated among signaling outgoing from PRG4^+^ lining fibroblasts in RA in comparison to PsA (Fig. 6d), OA and NIC (Extended Data Fig. 8d,e). Higher numbers of MHC-II interactions outgoing from the RA lining cluster in comparison to PsA were also evident in the spatial data (Extended Data Fig. 8f), with increased levels of MHC-II and CD74, in RA lining compared to PsA (Extended Data Fig. 8g,h). CD74 has a dual role acting as a receptor for migration inhibitory factor (MIF) on the cell surface and intracellularly facilitating antigen presentation on MHC-II^16,17^. In our scRNAseq dataset, in RA 72.3% of CD74-expressing fibroblasts in the sublining and 92.4% in the lining were also expressing MHC-II (Fig. 6e). These findings were confirmed on the protein level by multiplex fluorescence staining – 94.7% of CD74-expressing lining fibroblasts (characterized by CD55 expression) were also expressing MHC-II (Extended Data Fig. 8i,j) indicating a role of CD74 in antigen presentation in RA synovial fibroblasts. Flow cytometry analysis in murine models further confirmed elevated *CD74* expression in lining fibroblasts from RA model compared to PsA model (Fig. 6f, Extended Data Fig. 9). Also, sublining fibroblasts, in particular CXCL12^+^, CXCL14^+^ and CHI3L1^+^ synovial fibroblasts expressed high levels of MHC-II and *CD74* (Extended Data Fig. 8k) and MHC-II was also expressed in 3D micromass organoid in fibroblasts from both lining and sublining layer (Fig. 6g). However, in contrast to PRG4^+^ lining fibroblasts, enrichment of antigen processing and presentation pathways in sublining fibroblasts was not specific to RA (Extended Data Fig. 8l), suggesting that disease-specific mechanisms in the lining may involve interactions with FABP5⁺ iDC3s and SPP1⁺ macrophages, which are also localized to this compartment (Extended Data Fig. 8n).

RA PRG4^+^ lining fibroblast - but not other synovial fibroblast populations – exhibited specific enrichment of IFNγ response pathways (Fig. 6i, Extended Data Fig. 8m) which are known to enhance the antigen presentation pathway. Activity inference analysis further indicated up-regulation of the JAK-STAT signaling pathway, a downstream effector of IFNγ, in RA lining fibroblasts relative to other diagnoses (Fig. 6b). We confirmed this finding in lining synovial fibroblasts from a publicly available dataset^11^ (Extended Data Fig. 8a). MHC-II pathway enrichment in PRG4⁺ lining fibroblasts was observed across all synovial pathotypes but was most pronounced in the lympho-myeloid subtype (Fig. 6h), suggesting that IFNγ produced by infiltrating lymphocytes might stimulate antigen-presenting pathways in lining fibroblasts. However, *IFNG* expression in T cells and expression levels of IFNγ receptors (*IFNGR1*, *IFNGR2*) in RA PRG4⁺ fibroblasts were overall low and not elevated compared to PsA (Extended Data Fig. 8o,p). Moreover, cell–cell interaction analysis did not indicate enhanced IFNG–IFNGR signaling or increased signaling from T cells to fibroblasts in RA synovium.

To identify alternative upstream signals driving IFN response gene expression in RA lining fibroblasts, we performed ligand–receptor correlation analysis. This analysis indicated that IFN response–inducing signals originated predominantly from endothelial cells and fibroblasts (Extended Data Fig. 8q). Cell-cell interaction analysis showed that IL-6 signaling was among the top signals received by PRG4^+^ lining fibroblasts in RA in comparison to PsA (Extended Data Fig. 8r). *IL6* expression was higher in fibroblasts from synovium with lympho-myeloid pathotype (Fig. 6j) and *IL6R* expression was significantly higher in PRG4^+^ lining fibroblasts of RA patients in comparison to PsA (Extended Data Fig. 8s), suggesting a paracrine network of stromal and vascular cues contributing to IFN pathway activation in RA. Indeed, both IFNγ and IL-6 stimulations were able to induce HLA-II expression on synovial fibroblasts both on gene (Extended Data Fig.10a) and protein levels (Extended Data Fig. 10b,c). At the same time, stimulation of synovial fibroblasts with supernatants from activated T cells resulted in significant increase of IL-6 production, while stimulation with IFNγ alone failed to increase IL-6 concentration in fibroblast’s supernatants (Extended Data Fig. 10d). Analysis of synovial fibroblasts transcriptome showed that *IL6* was expressed predominantly by sublining fibroblasts (specifically by CXCL14^+^, CXCL12^+^ and CHI3L1^+^ SFs) and its expression was higher in RA in comparison to other diagnoses (Extended Data Fig. 10e).

To further investigate whether the expression of antigen-presentation and other IFNγ response pathway-related genes can be also induced by IL-6 stimulation in SFs we re-analyzed Tsuchiya *et al.* dataset^13^. Pathway analysis in synovial fibroblasts after IFNγ, IFNα or IL-6 stimulation showed significant up-regulation of both antigen presentation and IFNγ response pathways in each stimulation (Extended Data Fig. 10f,g), showing that IL-6 can induce transcriptional changes similar to those driven by IFN. Among the top genes up-regulated in PRG4^+^ fibroblasts in RA versus PsA in the antigen-presentation and IFNγ response pathways the majority (60% and 85.5% for antigen presentation and IFNγ response pathways) were overlapping with the genes up-regulated by both IFNγ and IL-6 stimulation (Extended Data Fig. 10h,i).

Inspired by the approach of Sinha *et al.*^18^, we applied a similar framework to predict the cytokine stimulation of lining synovial fibroblasts. For model training, we used the dataset from Tsuchiya *et al.*^13^, restricting the analysis to cells stimulated with either IL-6 or IFNγ. The classification identified on 78.2% of RA lining fibroblasts stimulated by IL-6, and only 19% by IFNγ (Fig. 6k, Extended Data Fig. 10j,k). Probability distributions differed between classes: IL6 scores were strongly concentrated near 1, reflecting confident predictions (Extended Data Fig. 10l). These findings further support the idea of IL-6 being the driving force of RA-specific synovial inflammation.

Based on these data, we hypothesize that increased levels of IL-6 production by sublining fibroblasts in RA induce an antigen-presenting phenotype in PRG4⁺ lining fibroblasts. Whether this increased antigen-presentation by synovial lining fibroblasts leads to activation of the adaptive immune system within the RA synovium remains to be determined. Nevertheless, given the aberrant production of CHI3L1 in RA synovium as well as previous data on arthrogenicity of CHI3L1 in mice, we aimed to see if CHI3L1 can be presented in HLA class II alleles associated with RA (shared epitope alleles^19^). Prediction of MHC-II peptide binding^20^ showed that higher number of peptides derived from the CHI3L1 protein can bind to MHC-II alleles associated with RA than to other MHC-II alleles, and such differences were not observed for other proteins of similar length (Fig. 6l, Extended Data Fig. 10m).

## Discussion

In our analysis, we demonstrate that increased interaction between fibroblasts and macrophages is a predominant characteristic of RA synovium. We identified MerTK^-^ SPP1^+^ macrophages as the main subtype of activated macrophages in RA, while several fibroblast subtypes were expanding in the RA synovium. We found a specific activation of synovial fibroblasts in the synovial lining layer in RA with enhanced antigen presentation pathways. Additionally, we identified a distinct CHI3L1-producing synovial fibroblast population that emerged specifically in RA synovium and might contribute to the initiation of an autoimmune response. Taken together, our data support the hypothesis that innate immune activation together with stromal activation, plays a central role in RA pathogenesis.

Our analysis focused on macrophage activation and fibroblast expansion in RA. In PsA, we observed increased fibroblast-fibroblast and endothelial-fibroblasts interactions. While in this analysis we did not reveal major differences in endothelial cells between RA and PsA, neither in histological features, cell proportions, nor differential gene expression, a more in-depth analysis, potentially involving further subclassification of endothelial cell types, may be necessary to uncover distinctive endothelial activation between RA and PsA.

Previous studies have highlighted the critical role of stromal–myeloid cell interactions in shaping the inflammatory environment in RA and showed that synovial fibroblasts can shape the phenotype of macrophages and vice versa^2,21,22^. Our analysis suggests that although signal transduction from macrophages to fibroblasts is prominent in RA synovium, it is not specific to RA but rather represents a general inflammatory feature of synovium. In contrast, signal transduction from the expanded and activated compartment of synovial fibroblasts to macrophages was an RA-specific feature. Similarly, the presence of MerTK^-^ SPP1^+^ macrophages has been identified in various inflamed tissues including RA at very early stages of disease^21^ and has been shown to correlate with disease activity^23^. However, based on our data their key marker gene, SPP1/osteopontin is specifically important in RA and serves as central signaling mediator in the interaction between MerTK^-^SPP1⁺ macrophages and other synovial cell populations in RA. SPP1/osteopontin has previously been implicated in promoting inflammatory and immune pathways. Our findings indicate that SPP1 may also contribute to the metabolic reprogramming of macrophages within the RA synovial microenvironment. Previously, it was shown that in vitro differentiated monocyte-derived macrophages from RA patients were metabolically changed^24,25^. Therefore, already circulating monocytes might be affected by the high serum SPP1 levels in RA patients. However, SPP1 levels in serum did not significantly differ between RA and PsA patients in our study, synovial fluid showed a trend toward higher levels in RA, arguing for local synovial activation. Interestingly, stimulation with SPP1 was previously shown to induce the expression of MMPs in synovial fibroblasts^26^. Therefore, increased production of SPP1 in RA synovium might also be a key event in the induction of invasive behavior of RA lining fibroblasts.

In the fibroblast populations, the most striking hallmarks of RA were the adaptive immune pathways active in lining synovial fibroblasts and the increased expression of CHI3L1 in sublining. Beyond their erosive capabilities, we hypothesize that the transformation of lining synovial fibroblasts in RA contributes to the breakdown of a well-defined lining structure, expanding into a more diffuse zone. This altered architecture may facilitate interactions between adaptive immune cells and specific myeloid populations, including MerTK^-^SPP1⁺ macrophages and activated FABP5⁺ iDC3 cells engaging with T cells^1^. The presentation of autoantigens by RA synovial fibroblasts within the lining niche appears plausible^27,28^ but requires further experimental validation. CHI3L1 (YKL-40/chondrex/HC gp-39) has been identified as a T cell autoantigen in the context of the HLA shared epitope. CHI3L1-derived peptides are presented by HLA-DR molecules and recognized by peripheral T cells from RA patients^7^. Moreover, in a murine model, these peptides were shown to induce arthritis, supporting their pathogenic relevance^8^. Intriguingly, we found a positive correlation of serum CHI3L1 levels with age, which might be connected to increased risk of autoimmunity with ageing.

The origin of the specific expansion and activation of the fibroblast populations in the lining as well as in the sublining remains elusive. Our data indicate that the IFN/IL-6/Jak-STAT signaling axis plays a central role in the activation of adaptive immune pathways in the lining in RA and in the induction of CHI3L1 expression by synovial fibroblasts. This activation of the IFN/IL-6/Jak-STAT signaling axis is particularly significant as CHI3L1 can, in turn, directly bind to the STAT3 coiled-coil domain^29^. This mechanism is strongly supported by observations within the RA synovium, where activated fibroblasts are characterized by the enrichment of STAT family transcription factor motifs in their open chromatin and the widespread presence of phosphorylated STAT1, confirming that this pathway is a central driver of their pathogenic phenotype^30^. The epigenetic imprinting of the CHI3L1 locus in synovial fibroblasts could reflect an early, preclinical priming event involving the Jak-STAT pathway, such as a viral infection. In combination with the strong genetic predisposition conferred by MHC class II variants, such an environmental trigger could promote a tissue-specific response characteristic of RA. An investigation of peripheral blood cells from patients with early RA also suggested a role for interferon-regulated epigenetic programming in the breach of tolerance in early stages of the disease in peripheral blood^31^.

We have to acknowledge that our study is largely based on dissecting observed differences in cell activation between RA and other arthritides and does not provide a causal explanation for these differences. Nevertheless, we think that our data accurately captured unique characteristics of RA synovium and provides important novel clues on how activation of the adaptive immune system could be driven (via production of auto-antigens) and supported (via providing a tissue niche) by fibroblast activation in a disease-specific manner.

## Methods

### Ethical statement

The collection and experimental usage of the human samples was approved by the ethical commission of the Kanton Zurich (swissethics numbers: 2022-01818, 2019-00674, 2016-02014, 2021-00092 and 2019-00115). Informed consent was obtained from all patients and healthy volunteers. All experiments have been performed in accordance with international and institutional ethical guidelines.

Mouse experiments were approved by the Institutional Committee of Protocol Evaluation in conjunction with the Veterinary Service Management of the Hellenic Republic Prefecture of Attika, according to all current European and national legislation, and were performed in accordance with relevant guidelines and regulations, under the relevant animal protocol licenses with number 488046-26/05/2022.

### Synovial tissue collection and processing

Synovial biopsies were obtained from patients with active RA, PsA and UA by ultrasound-guided fine-needle biopsy after obtaining informed consent. All patients fulfilled the ACR/EULAR classification criteria^35^ for RA and CASPAR criteria for PsA^36^. UA did not fulfill criteria for any classified chronic arthritis. From two RA patients, two biopsies from two different joints were included. Synovial tissues of OA patients were obtained from joint replacement surgeries. Non-inflammatory control synovial tissues were obtained from arthroscopic surgeries for meniscopathy, absence of synovitis was confirmed histologically. All synovial tissues used in this study were processed fresh, and the presence of synovium was confirmed histologically for all included samples. Synovial tissues were digested as previously described in Edalat *et al.*^37^. Briefly, synovial tissues were minced and digested in a mixed enzymatic-mechanical dissociation including 100 mg/mL Liberase TL (Roche) and 100 mg/mL DNAse I (Roche); red blood cells were lysed (Red Blood Lysis buffer, Roche), and synovial cell pellets washed. Cells were washed and counted on a LUNA automated cell counter (Logos Biosystems). Synovial single-cell libraries were prepared using the 3’ v3.0 and 3’ v3.1 protocols and reagents from 10X Genomics according to the manufacturer’s instructions targeting the encapsulation of up to 6000 cells. The quality and quantity of cDNA and libraries were assessed with Agilent Bioanalyzer (Agilent Technologies). Diluted 10nM libraries were pooled in equimolar ratios, and the library pools were sequenced on the Illumina NovaSeq6000 sequencer (paired-end reads, R1 = 28, i7 = 8, R2 = 91) at Functional Genomics Center Zurich, the University of Zurich and ETH Zurich, Switzerland.

### Single-cell RNA sequencing (scRNAseq) analysis

In addition to in-house generated scRNAseq dataset, publicly available scRNAseq datasets from national center for biotechnology information (NCBI) ascension number GSE200815 (from Floudas *et al.*^11^) and BioStudies E-MTAB-14339 (from Raimondo *et al.*^38^) were used. Transcript count tables were generated from fastq files using cellranger count (cellranger, version 6.0.0, 10x Genomics) and the reference Genome GRCh38 (GENCODE v32/Ensembl 98). The filtered count matrix was used for downstream analysis. The analysis was performed in R (version 4.0.3 and version 4.4.2). Seurat^39^ (version 5.2.1) was used for the scRNAseq data analysis. The preprocessing consisted of the following steps: ambient RNA removal using SoupX^40^ (version 1.6.2), doublet detection using DoubletFinder^41^ (version 2.0.6) and scDblFinder^42^ (version 1.20.2), cell filtering using thresholds set manually per sample (thresholds for nFeature_RNA, nCount_RNA, percent.mt), normalization (log-transformed normalized expression values) using Seurat, 2000 highly variable genes selection using Seurat, data scaling using Seurat, PCA dimensionality reduction using Seurat (number of PCs for downstream analysis was selected manually using ElbowPlot). For major clusters annotation unintegrated data were used, neighbors and clusters were identified and Uniform Manifold Approximation and Projection (UMAP) was calculated using Seurat. Major cell types were annotated manually. Marker genes were determined with Wilcoxon tests. For finer subpopulation annotation per cell type (fibroblasts, myeloid cells, and dendritic cells) the data were integrated across synovial biopsies using STACAS^43^ (standard integration) (version 2.3.0). Cell subpopulation annotation followed the same steps as the main cell type annotation, starting with highly variable gene selection for each cell subset. Proportions of cell populations and subpopulations were calculated and analyzed using differential composition analysis (DCATS^34^ version 1.4.0) and a permutation test (scProportionTest^44^ version 0.0.0.9000). Differential gene expression analysis was conducted using the FindMarkers function in Seurat, which implements the MAST^45^ framework (version 1.32.0). Sample identifiers were included as covariates to account for inter-sample variability. Pathway analysis was performed using clusterProfiler^46^ (version 4.14.6). Cell-cell communication analysis was performed using CellChat^33^ (version 2.2.0) and multinichenetr^32^ (version 2.1.0). Gene signature scores were computed using UCell^47^ (version 2.10.1). Python (version 3.12) implementation of decoupler^9^ (version 2.1.1) was used for pathway activity inference analysis (PROGENy).

### Spatial transcriptomics

Synovial tissues were fixed with 4% paraformaldehyde and embedded in paraffin. The quality of RNA was assessed with Agilent Bioanalyzer (Agilent Technologies) via calculating DV200. Formalin-fixed paraffin-embedded (FFPE) synovial tissues with DV200 > 90% were used. The FFPE blocks were sectioned (5-μm sections), stained with H&E, permeabilized at 37 °C for 12 min and polyadenylated mRNA was captured by primers bound to the slide. Reverse transcription, second strand synthesis, cDNA amplification and library preparation proceeded using the Visium Spatial Gene Expression Reagent Kits for FFPE (10x Genomics CG000407) according to the manufacturer’s protocol. After evaluation by real-time PCR, sequencing libraries were prepared with 17-20 cycles of PCR. Indexed libraries were pooled equimolar and sequenced on a NovaSeq X Plus (Illumina) (paired-end 150 bp) at Functional Genomics Center Zurich, the University of Zurich and ETH Zurich, Switzerland.

### Spatial transcriptomics data analysis

Spatial sequencing data from the synovial tissue were aligned using the Space Ranger (v.1.2.2, 10x Genomics) pipeline to the reference Genome GRCh38 (GENCODE v32/Ensembl 98). The analysis was performed in R (version 4.4.3). Seurat^39^ (version 5.3.0) was used for the analysis. The preprocessing was performed per sample. Genes expressed in fewer than 10 spatial spots were excluded from downstream analyses. Spots with low numbers of detected genes were also excluded, with filtering thresholds set manually for each sample. Spots with fewer than 200 total detected counts (nCount_Spatial) were removed across all samples. Each dataset was normalized and variance-stabilized using Seurat’s SCTransform method. The resulting corrected values were used for downstream integration with Harmony^48^ (version 1.2.3). Neighbors and clusters were identified and UMAP was calculated using Seurat. The clusters were annotated manually. Marker genes were determined with Wilcoxon tests. Spatial transcriptomics data were deconvoluted using Spacexr^49^ (version 2.2.1) by integrating it with scRNAseq datasets to estimate cell type proportions at each spatial location. Cell-cell communication analysis was performed using CellChat^33^ (version 2.2.0). Gene signature scores were computed using UCell^47^ (version 2.10.1).

An independent analysis was performed using the STutility^50^ (version 0.1.0) to explore spatial neighbourhood relationships. Raw Visium data were imported and filtered, applying thresholds of 100 minimum UMI counts per gene and 500 per spot. Regional neighbour relationships were identified using the RegionNeighbours function.

### PBMC scRNAseq

Cryopreserved peripheral blood mononuclear cells (PBMCs) were thawed rapidly at 37 °C, transferred to prewarmed RPMI supplemented with 10% FCS, and washed twice to remove residual DMSO. Cells were resuspended in PBS containing 0.04% BSA, and viability was assessed using a LUNA automated cell counter (Logos Biosystems). Following centrifugation, cell pellets were treated with freshly prepared DNase I solution on ice to eliminate extracellular nucleic acids, washed, and resuspended in PBS/BSA. For nuclei isolation, approximately 1 × 10⁵ cells per sample were lysed on ice in buffer containing 0.1% Tween-20, 0.1% NP-40 substitute, 0.01% digitonin, and 1% BSA, followed by gentle washing in RNase inhibitor–containing buffer. Nuclei quality and concentration were determined by trypan blue staining, and samples were adjusted to 4,000 nuclei/μL. Nuclei were processed immediately using the Chromium Next GEM Single Cell Multiome ATAC + Gene Expression platform (10x Genomics) according to the manufacturer’s protocol, targeting a recovery of 6,000 nuclei per sample. GEM generation, barcoding, cDNA and ATAC library construction, and amplification were performed following 10x Genomics instructions. Library size and concentration were assessed using an Agilent Bioanalyzer (Agilent Technologies), and pooled libraries were sequenced on an NovaSeq 6000 (Illumina) (paired-end reads, R1 = 28, i7 = 8, R2 = 91) at Functional Genomics Center Zurich, the University of Zurich and ETH Zurich, Switzerland.

PBMC scRNAseq data were processed following the workflow described above. Briefly, doublets were removed using scDblFinder^42^, ambient RNA contamination was corrected with SoupX^40^, low-quality cells were filtered via manual quality control, and multiple datasets were integrated using STACAS^43^. Cell types were annotated manually. Myeloid cell and DC populations were integrated with corresponding synovial subpopulations using STACAS^43^.

### Monocyte Isolation and Differentiation to MDMs

Peripheral blood, freshly extracted from healthy volunteers, was diluted with 1X PBS at a ratio 1:1. PBMC isolation was then performed using Histoplaque (Sigma-Aldrich). Monocyte isolation was performed using positive selection with CD14 Microbeads (Miltenyi Biotec) following the company’s isolation protocol. Briefly, PBMCs were washed with MACS Buffer (PBS, 0.5% bovine BSA, 2 mM EDTA) and centrifuged at 300g for 10 min. The supernatant was aspirated completely, and the cell pellet was resuspended in 80 μL of MACS buffer. 20 μL of CD14 MicroBeads was added and mixed well with the cell suspension. The cell suspension was incubated for 15 min in the refrigerator (+2 to +8°C). Following incubation cells were washed by adding 1−2 mL of buffer and centrifuged at 300g for 10 min. The supernatant was discarded, and the cell pellet was resuspended in 1 mL of MACS Buffer. Positive cell sorting was performed at the AutoMACS® NEO separator (Miltenyi Biotec). After separation, monocytes were placed in 12-well plates (at a density of 5 × 10^5^ cells per well) and GM-CSF (PeproTech; 15 ng/mL) was added and replaced every 48h for seven days.

### ELISA

The levels of cytokines, matrix metalloproteinases (MMPs), pro-collagen type I alpha 1, chitinase-3-like protein 1 (CHI3L1), and osteopontin (SPP1) were evaluated using ELISA kits according to the manufacturer’s protocol. Detailed information on ELISA kits is provided in Supplementary Table Reagents. Optical density was measured at 450 nm with wavelength correction at 540 nm using a multimode microplate reader (BioTek Synergy H1 Multimode Reader, Agilent). Protein concentrations were calculated using a four-parameter logistic standard curve. All samples were analyzed in duplicates.

### Lactate Assay

Lactate levels in cell culture supernatants were measured using the Lactate-Glo™ Assay (Promega) following the manufacturer’s instructions. Supernatants from cultured synovial fibroblasts were diluted 1:100, and those from MDMs were diluted 1:50 in PBS. 50 μL of each diluted sample was mixed with 50 μL of Lactate Detection Reagent in white opaque 96-well plates and incubated for 60 minutes at room temperature in the dark. Luminescence was measured using a BioTek Synergy H1 plate reader (0.5-second integration time, AutoGain, 6.5 mm read height; Agilent). Raw luminescence values were background-corrected by subtracting the signal from media-only control wells.

### Mitochondrial Membrane Potential Assay

A working solution of Image-iT™ TMRM Reagent (Invitrogen) at 100 nM in complete DMEM (Gibco, Life Technologies) supplemented with 10% fetal calf serum, 2 mM glutamine, 50 U/mL penicillin, 50 U/mL streptomycin, 10 mM HEPES, and 0.2% amphotericin B (Gibco) was prepared and applied to the cells, followed by incubation for 30 min at 37°C. Nuclear counterstaining was performed using Hoechst 33342 (Thermo Fisher Scientific). After staining, cells were washed and replaced with fresh complete medium. Fluorescent images were acquired using the Thermo Scientific™ CellInsight CX7 High Content Screening Platform. Image analysis and feature extraction were performed using CellProfiler software (version 4.2.1).

### SF Isolation and Stimulation

#### Cell culture

Synovial tissues were digested using Accutase and Liberase at 37°C for 1 hour, and the resulting SFs were cultured in Dulbecco’s Modified Eagle Medium (DMEM; Life Technologies). The culture medium was supplemented with 10% fetal calf serum (FCS), 50 U/ml penicillin/streptomycin, 2 mM L-glutamine, 10 mM HEPES, and 0.2% amphotericin B (all from Gibco). Cell cultures were tested and confirmed to be free from mycoplasma contamination using the MycoAlert mycoplasma detection kit (Lonza). SFs from passages 4 to 6 were used for experiments.

#### Synovial Fibroblast *in vitro* stimulation

SFs derived from RA, OA, and NIC were stimulated with human recombinant TNFα (10 ng/mL; R&D Systems), human recombinant IL-1β (1 ng/mL; R&D Systems), human recombinant IL-6 (50ng/mL; R&D Systems), human recombinant IFNγ(10 ng/mL; ImmunoTools), human recombinant CHI3L1 (2 μg/mL; R&D Systems) or human recombinant TGFβ (10ng/mL; PeproTech, ThermoFisher) for 48 h.

### RNA Isolation, cDNA Synthesis, and RT-qPCR

Total RNA was isolated using the miRNeasy Mini Kit (Qiagen) with on-column DNase I digestion. cDNA was synthesized using MultiScribe Reverse Transcriptase with random hexamers (Thermo Fisher Scientific) according to the manufacturer’s instructions. Quantitative PCR was performed on a 7900HT Fast Real-Time PCR System (Thermo Fisher) using FastStart Universal SYBR Green Master Mix (Roche). No template control samples, dissociation curves and samples containing the untranscribed RNA were measured in parallel as controls. Gene expression levels were analyzed using the comparative CT method (2⁻ΔCT or 2⁻ΔΔCT) and are presented as log2 fold changes relative to the control condition. *RPLP0* served as the reference gene for normalization. Specific primers used are listed in Supplementary Table Sequences.

### Immunohistochemistry (IHC) Staining of CHI3L1 in RA synovial tissue

FFPE rheumatoid arthritis (RA) synovial tissue blocks were sectioned at 3 μm thickness using a HistoCore Autocut microtome (Leica Biosystems) and mounted onto Superfrost® Plus Menzel glass slides (Thermo Fisher Scientific). Histological staining with hematoxylin and eosin (H&E) was performed according to standard protocols to assess tissue morphology. Immunohistochemistry staining for CHI3L1 was performed on a BOND-MAX instrument (Leica Biosystems) using the BOND Polymer Refine Detection kit (Leica Biosystems) according to manufacturer’s instructions. Briefly, sections were deparaffinized, followed by heat-induced epitope retrieval (HIER) with citrate-based (pH 6, BOND Epitope Retrieval Solution 2, Leica Biosystems)) or ethylenediaminetetraacetic acid (ETDA)-based buffer (pH 9, BOND Epitope Retrieval Solution 2, Leica Biosystems). Next, sections were blocked of unspecific antibody binding, followed by incubation with primary and secondary antibodies. 3,3’-Diaminobenzidine (DAB) was used as an antibody detection chromogen and counterstaining was performed with hematoxylin. Detailed information on antibodies used for IHC is provided in Supplementary Table Reagents.

Whole slide scans of IHC stained synovial tissue sections were semi-automatically imaged on an AxioScan.Z1 (Zeiss) microscope scanner at 20X magnification.

### Multiplex Immunofluorescent Staining

High-plex spatial protein profiling was performed on formalin-fixed, paraffin-embedded (FFPE) synovial tissue sections using the fully automated COMET™ platform (Lunaphore). Slides were incubated for 1 h at 60 °C, manually deparaffinized with xylene (2 × 5 min) and 100% ethanol (2 × 3 min) and subjected to heat-induced antigen retrieval in a PT module (Erpedia) at 99 °C. Sections were then blocked with 5% horse serum (Vector Laboratories, S2000-20).

Sequential immunofluorescence (seqIF™) was executed automatically across 18 cycles of primary antibody incubation, secondary antibody detection, imaging, and rapid antibody elution. A customized panel of 31 validated primary antibodies was used, with signal detection via TRITC-, Cy5-, and Cy7-conjugated secondary antibodies. Detailed antibody information is provided in Supplementary Table Reagents.

Following image acquisition, hyperplex images were processed using Horizon™ software (version 2.4.0.0 Lunaphore, which performed image registration, background subtraction, and cell segmentation for quantitative analysis. Cell segmentation employed a deep learning–based algorithm derived from InstanSeg, optimized for COMET data, and cell classification was performed by manual feature thresholding.

### DNA Methylation Analysis and ChIP-Seq Tracks

For DNA methylation analysis, data from the cohort described in Karouzakis *et al.*^51^, along with a separate unpublished cohort profiled using Illumina 450k and EPIC microarray chips, were integrated in R (version 4.4.2) using the minfi package^52^ (version 1.50.0). The ssNoob normalization method was applied to obtain β-values for CpG sites. For ChIP-seq, publicly available data (GEO accession - GSE163548) from Ge *et al.*^53^ were used. BAM files were normalized using the Reads Per Genomic Content (RPGC) method with the bamCoverage function from deepTools^54^ (version 3.5.6), and subsequently converted to bigWig format. All tracks, including DNA methylation and histone modifications (H3K4me3 and H3K27ac), were visualized using Gviz^55^ package (version 1.48.0).

### Silencing

SFs were seeded at 1.5 × 10⁵ cells per well in 6-well plates and transfected with 50 nM antisense LNA GapmeR targeting *CHI3L1* (5’ - CGAGGATTCTATGGAC - 3’, Qiagen). Transfections were performed using Lipofectamine 2000 (Invitrogen) according to the manufacturer’s instructions. A non-targeting LNA GapmeR (Negative Control A, Qiagen) was included as a transfection control. Total RNA was extracted 48 h after transfection and knockdown efficiency was assessed by quantitative RT-PCR.

### Bulk RNAseq

Raw sequencing reads were quality-checked and trimmed using Trim Galore (Babraham Bioinformatics) to remove adapters and low-quality bases. Trimmed reads were aligned to the human reference genome (hg38) using STAR^56^. Gene-level counts were summarized with featureCounts^57^. Count data were imported and filtered to remove low-count genes. Data were normalized and variance-stabilized using DESeq2^58^ (version 1.46.0). Principal component analysis (PCA) was used to assess sample clustering. Differential expression analysis was performed with DESeq2^58^, and gene set enrichment analysis (GSEA) was conducted using clusterProfiler^46^ (version 4.14.6) on ranked log2 fold changes. Sample “492” was excluded from downstream silencing RNA-seq analyses owing to insufficient CHI3L1 knockdown efficiency.

### Stimulation of SFs with T Cells Supernatants

Peripheral blood, freshly extracted from healthy donors and RA patients, was diluted with 1X PBS at a ratio of 1:1.5. PBMC isolation was then carried out using SepMate tubes (Stemcell technologies) with 14 mL Histoplaque (Sigma-Aldrich). Red blood cell lysis was performed using 4 mL of 1X Red blood cell lysis buffer (BD Biosciences). PBMCs were washed with PBS, centrifuged at 300g for 5 min at 4°C and counted. Pan T cell magnetic purification was performed using autoMACS (Miltenyi Biotec) and pan T cell isolation kit (Miltenyi Biotec) following the company’s isolation protocol. Pan T cells were counted and placed in 24-well plates with RPMI medium (1% NEAA, 1% Sodium Pyruvate, 1% Glutamax, 1% Pen/strep, 10% FBS and 50uM b-mercaptoethanol) containing Immunocult Human CD3/CD28 T Cell Activator (Stemcell technologies, #10971) and IL-2 (dilution 1:500, Roche). After 24 h Pan T cells were collected, centrifuged, and placed in fresh RPMI medium containing only IL-2 at a concentration of 1.5-2 million cells/mL. After 48 h the supernatants were collected and used for the synovial fibroblast activation. After 48 h of fibroblast activation, the supernatants were collected for ELISA.

### THP-1 Stimulation with rhCHI3L1

THP-1 (human monocytic cell line, TIB-202) were obtained from the American Type Culture Collection (ATCC). Cells were grown in RPMI 1640 medium with L-glutamine and 25 mM HEPES buffer supplemented with 10% fetal bovine serum, 100 U/mL penicillin, and 100 mg/mL streptomycin (Gibco) at 37°C in a humidified atmosphere containing 5% CO2 in air. THP-1 cells were differentiated into macrophages after incubation with 100 ng/mL phorbol-12-myristate-13-acetate (PMA; Sigma-Aldrich) for 24 h. They were then incubated with rhCHI3L1 (500 ng/mL and 2000 ng/mL, R&D Systems, #2599-CH) and/or LPS (2μg/mL, abbexa) for 24 h.

### Flow Cytometry Analysis of SFs

Flow cytometry analysis was performed using a BD FACS LSRFortessa cell analyzer equipped with five lasers and DIVA software (BD Biosciences). Cells were harvested, washed with PBS, and resuspended in FACS buffer (PBS containing 2% FBS and 1 mM EDTA). Cells were stained with eFVD Live/Dead Fixable cell viability marker 405 nm (Thermo Fisher Scientific) for 30 min at 4°C. Cells were washed with FACS buffer and stained with fluorochrome-conjugated antibodies against specific surface markers (HLA-DR, CD74, and CD45) for 30 min at 4 °C in the dark. Detailed antibody information is provided in Supplementary Table Reagents. Data were acquired with at least 10,000 events recorded per sample. FMOs were prepared for each fluorochrome. Analysis was performed using FlowJo software (version 10.09 and 10.10.0, TreeStar).

### Fluorescence-activated cell sorting (FACS), scRNAseq data processing and analysis of murine synovial tissue

Isolation of CD45^-^, CD31^-^, Pdpn^+^ synovial cells from A20^Znf7-KI^ mice (PsA-like arthritis)^59,60^ before onset (week 5) and established disease (week 25) was performed as previously described^61^. Synovial single-cell libraries were prepared using the 3’ v3.1 protocols and reagents from 10X Genomics according to the manufacturer’s instructions preparations. Libraries were sequenced at a Hiseq X sequencer (2x150bp / 700M reads total) (Macrogen).

Raw scRNAseq data were processed using Cell Ranger v7.1.0 (10x Genomics) with the mouse reference genome refdata-gex-mm10-2020-A to perform read mapping, UMI quantification, and generation of gene-by-cell count matrices. Downstream analysis was performed in R using Seurat^39^ version 5. Cells were filtered based on quality-control thresholds: cells with 1,000 - 7,000 detected genes (nFeature_RNA) and <10% mitochondrial content were retained for downstream analysis. Filtered data were normalized using SCTransform with the glmGamPoi method for variance stabilization, regressing out the effect of mitochondrial content, and the 3,000 most variable genes were selected for downstream dimensionality reduction. Principal component analysis (PCA) was applied to the scaled expression matrix, and the first 50 resulting components were used to construct a shared nearest-neighbor (SNN) graph for graph-based clustering (Louvain, with resolution = 0.8). The low-dimensional embedding was visualized using UMAP. To focus on SFs, clusters expressing canonical SF marker genes were identified and retained for all subsequent analyses.

For the analysis of SFs derived from RA-like arthritis we employed the published scRNAseq datasets (BioProject under accession code PRJNA778928).

### Flow cytometric analysis of murine CD74 expression

Freshly isolated SFs from hTNF-Tg (RA-like arthritis), A20^Znf7-KI^ mice (PsA-like arthritis) and wild type (WT) mice were processed for the detection of CD74 expression. Upon washing the cells with PBS containing DNase, they were blocked in 1% BSA in PBS and Fc blocker (unlabelled anti-CD16/32, Biolegend 101302) for 10 min at 4 °C and stained with fluorophore-conjugated antibodies for 20 min at 4 °C. Detailed antibody information is provided in Supplementary Table Reagents. After washing with PBS, cells were resuspended in FACS buffer (PBS, 1%BSA). For each sample, an FMO control sample was analyzed, stained with all antibodies except for CD74, allowing us to determine the true background signal. The analysis was performed with BD FACSCanto II and the BD FACSDiva software and dead cells were excluded by Propidium Iodide staining. Analysis of the results were performed employing FlowJo software (v.10).

### Micromasses

Micromass organoid culture was used as a 3D cell formation as previously described by Kiener *et al.*^62^. Briefly, SFs from passages 3 to 10 from RA patients and human umbilical vein endothelial cells (HUVECs) (Lonza), were prepared for 3D culture. The cells were detached using trypsin (Gibco), washed with calcium-free PBS, and counted. Monocytes were isolated from blood of healthy donors and RA patients, using AutoMACS® NEO separator (Miltenyi Biotec) and counted. The cells were then suspended in cold Matrigel Matrix (Corning) at the following concentrations: FLSs: 2 × 10⁶ cells, HUVECs: 2 × 10⁶ cells, monocytes: 1 × 10⁶ cells. Droplets (35 μL) of the cell suspensions were placed onto ultra-low attachment culture dishes (Nuclon Sphera 12 well, Life Technologies 174931). The Matrigel was allowed to solidify for 45 min at 37°C. After gelation, the cultures were covered with EGM-2 medium (Lonza) containing all supplements except hydrocortisone. The 3D cultures were maintained for 14 days, with medium changes every 2 days.

Micromasses were collected and immediately fixed in BD Cytofix™ (diluted 1:4 in DPBS, BD Biosciences) for 20 hours at 4°C. Following fixation, micromasses were washed twice with washing buffer containing 0.05% sodium azide in PBS (Sigma-Aldrich) and stored at 4°C in the same buffer.

For immunostaining, fixed micromasses were embedded in 4% low-melting agarose (Sigma-Aldrich) and sectioned into 70-100 μm-thick sections using a vibratome (Leica VT1200S) equipped with conventional razor blades. The sections were then washed for 20 min at room temperature on a shaker in washing buffer (dH*2*O supplemented with 0.1% Tris (Thermo Fisher), 1% mouse serum (Jackson ImmunoResearch) and 1% salmon sperm DNA (Thermo Fisher)).

Sections were then stained with 300 μL antibody staining mixture (see detailed antibody information in Supplementary Table Reagents) per section overnight at room temperature in the dark. The staining mixture was prepared in dH*2*O containing 0.1% Tris, 0.3% Triton-X100 (Bio-Rad), 1% Mouse Serum and 1% ssDNA to enhance permeabilization and reduce background. Following overnight incubation, tumor sections were washed twice on the shaker (10 min per wash). Finally, stained sections were mounted onto glass slides (Epredia) using an ImmEdge Pen (vector laboratories) and Fluoromount-G™ mounting medium (Thermo Fisher), then cover-slipped (Roth) for confocal imaging.

Images were acquired using a Stellaris 5 inverted confocal microscope (Leica Microsystems) equipped with a 40X oil immersion objective with a numerical aperture of 1.3. The confocal pinhole was set to 2.00 AU, resulting in an optical section thickness of 1.038 μm. Images were captured at a resolution of 1024x1024 pixels and 400 Hz, with a z-spacing of 1.5 μm. Tiled image stacks were acquired and fluorophore spillover was calculated and subtracted using Leica LAS X software. Image visualization was performed using Imaris software (9.7.2. and 10.1.1; Bitplane).

### Classification of lining fibroblasts by stimulation

Inspired by the approach of Sinha *et al.*^18^, a similar framework to predict the cytokine stimulation (IL-6 or IFNγ) of lining SFs was applied.

The classification was performed using R (version 4.5.0). For model training, dataset from Tsuchiya *et al.*^13^, consisting of bulk RNA sequencing data from fibroblasts derived from OA and RA patients, was used. The analysis was restricted to cells stimulated with either IL-6 or IFNγ, retaining the genes with at least 100 reads across samples.

Classification was carried out using the scPred^63^ package (version 1.9.2). scPred first performs dimensionality reduction and defines a predictive feature space by selecting principal components that are informative for the stimulation labels. The classification step relies on a support vector machine (SVM) with a radial basis function kernel. SVMs are supervised learning algorithms that identify an optimal separating hyperplane between class labels in a high-dimensional feature space. The model outputs class probabilities for each stimulation condition. Cells were assigned to a class if the corresponding probability exceeded the threshold of 0.5.

This threshold was validated on a test subsample from the Tsuchiya dataset, yielding high performance for IFNγ classification with a sensitivity of 0.982 and a specificity of 0.983, thus justifying the choice of 0.5.

After normalization of our single-cell data, it was projected into the trained feature space. The intersection of informative features between the training and test datasets included 1,839 of the 2,000 genes selected during training. Probability distributions differed between classes: IL-6 scores were strongly concentrated near 1, reflecting confident predictions, whereas IFNγ scores were generally lower and clustered around 0.6, suggesting greater classification uncertainty.

Several limitations should be mentioned. First, the model was trained on bulk RNA-seq data but applied to single-cell data, which may introduce biases. Second, training and testing were conducted on datasets from different sources, which could reduce generalizability. While the classifier showed strong internal performance, further validation is needed to confirm robustness in independent single-cell datasets.

### MHC-II epitope prediction

Potential HLA class II epitopes were predicted using NetMHCIIpan-4.0 (DTU Health Tech) with default settings. Protein sequences were fragmented into overlapping 14–23-mer peptides (sliding window of 1 amino acid). Peptide–allele pairs with a NetMHCIIpan %Rank EL ≤10 were classified as binders. To summarize by allele group, counts were aggregated across alleles belonging to the shared-epitope (SE) group versus the non-SE group (SE list: DRB1*04:01, *04:09, *04:13, *04:16, *04:21, *14:19, *14:21, *01:01, *01:02, *01:05, *04:04, *04:05, *04:08, *04:10, *04:19, *14:02, *14:06, *14:09, *14:13, *14:17, *14:20, 10:01; not-SE list: DRB101:03, *03:01, *03:05, *04:02, *04:03, *07:01, *08:01, *08:03, *09:01, *11:01, *11:04, *12:01, *13:01, *13:02, 13:03, 14:01, 14:54, 15:01, 15:03, 16:01, DRB301:01, DRB302:02, DRB401:01, DRB401:03, DRB501:01, DRB502:02). Allele grouping was performed based on published SE definitions^19^.

## Data availability

RNAseq and scRNAseq data generated in this study will be available upon manuscript acceptance from ArrayExpress.

Spatial transcriptomics data generated in this study will be available upon manuscript acceptance from ArrayExpress.

The mouse data analyzed in this study are part of ongoing work by the group of M.A. and are not publicly available. Access may be requested from the original authors, subject to their approval.

## Acknowledgements

We would like to thank Peter Kunzler, and other members of the laboratories of C.O. for helpful discussions and technical help.

We thank Dr. Miriam Marks for her assistance with sample collection. We thank Penelope Timpert for valuable input as a patient representative and for providing perspectives that helped shape the interpretation and contextualization of this study.

We thank the Functional Genomic Center Zürich for technical support and expertise. The authors would like to thank patients and institutions who participated in SCQM (participating institutions are listed on www.scqm.ch/institutions). The SCQM is supported by sponsors (www.scqm.ch/sponsors).

The authors used ChatGPT (GPT-5, OpenAI), Microsoft Copilot, and DeepL to assist in refining the language of the manuscript. The authors take full responsibility for the content.

This work was supported by the Foundation for Research in Rheumatology (FOREUM), Bangerter-Rhyner Stiftung and the Swiss National Science Foundation.

