## Supplementary data for "Disease-specific fibroblast-myeloid interactions in rheumatoid arthritis synovium"

#### Extended data

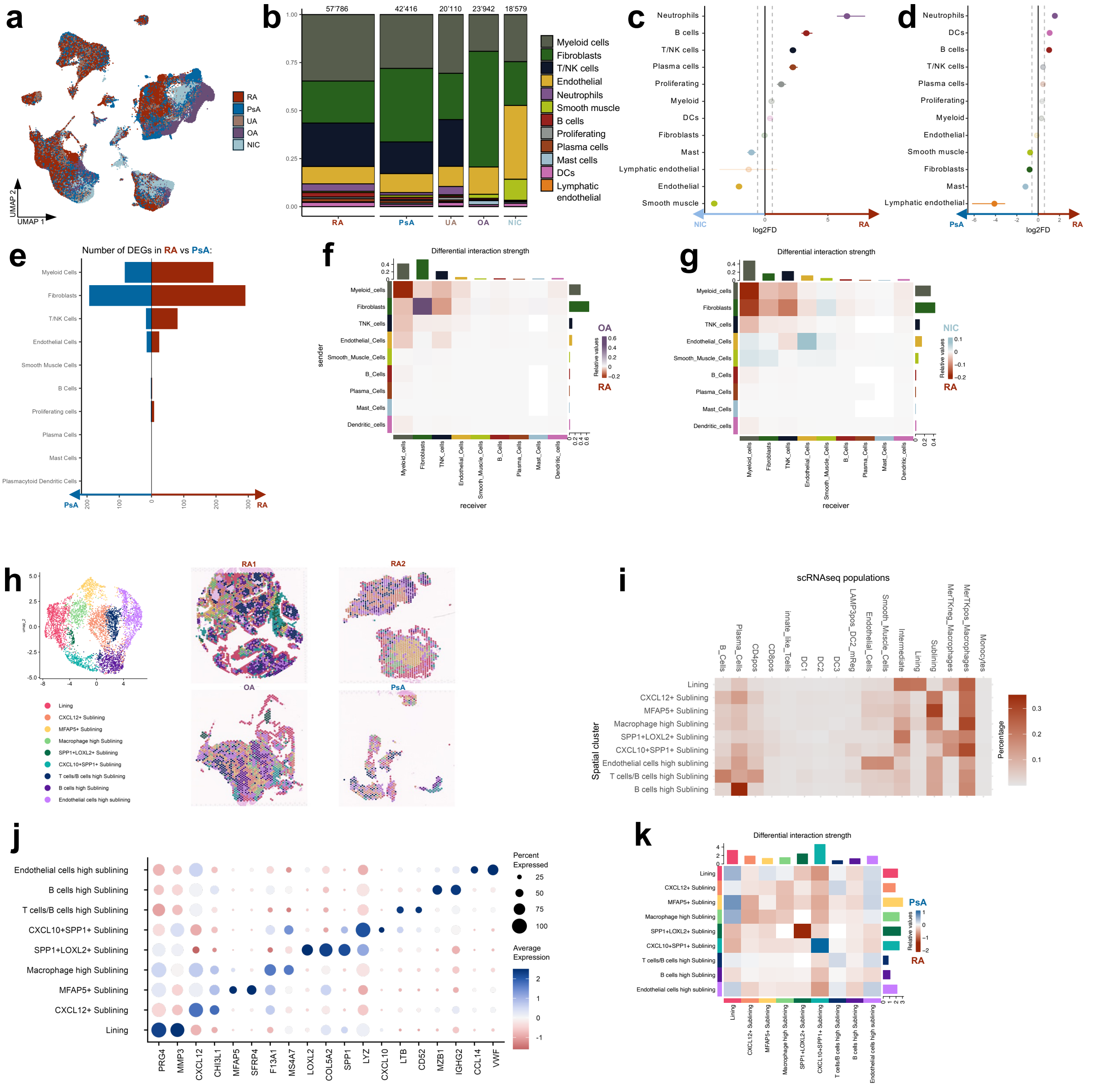


#### Extended Data Figure 1.

**a** UMAP of main synovial cell populations colored by diagnosis in unintegrated merged scRNAseq data. **b** Barplot of the proportions of the main cell populations across the diagnoses. The bar's width corresponds to the total number of cells per diagnosis (indicated above). **c** Dot plot of log2 fold differences in proportion of cell populations between RA and PsA calculated by permutation test (scProportionTest[44]). **d** Dot plot of log2 fold differences in proportion of cell populations between RA and NIC calculated by permutation test (scProportionTest[44]). **e** Number of DEGs (log2FC > 0.5 and p-value adjusted < 0.05) between RA and PsA by cell subpopulation. **f** Heatmap of differential interaction strength in OA versus RA according to CellChat[33]. **g** Heatmap of differential interaction strength in NIC versus RA according to CellChat[33]. **h** UMAP and spatial dim plots of spatial dataset clusters. **i** Heatmap of the percentage of scRNAseq subpopulations identified in the spatial dataset clusters by deconvolution analysis (using Spacexr[49]). **j** Bubble plot of spatial dataset clusters marker genes. **k** Heatmap of differential interaction strength in PsA versus RA according to CellChat[33] in spatial dataset.

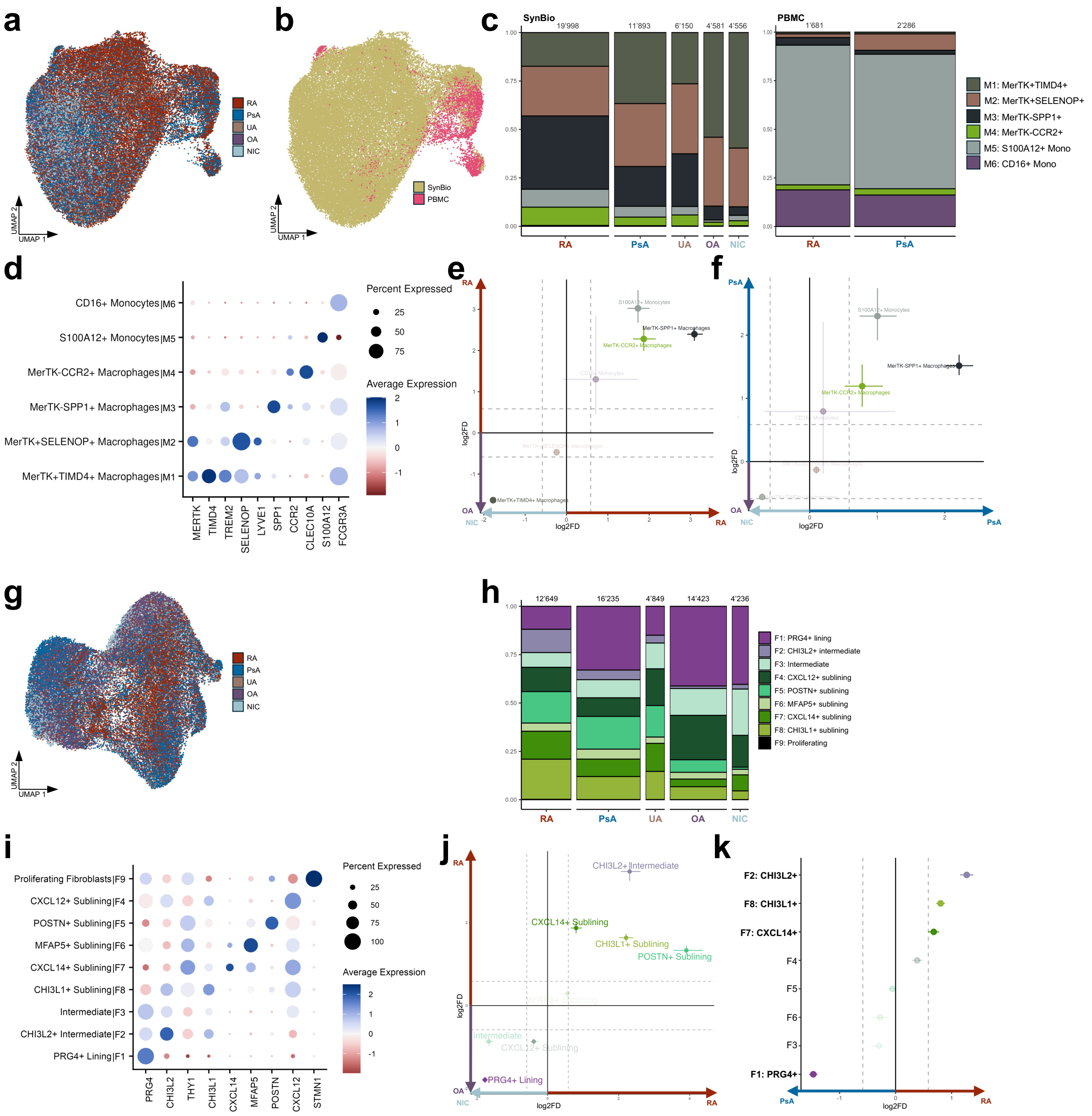


#### Extended Data Figure 2.

**a** UMAP of myeloid cell populations colored by diagnosis in integrated scRNAseq data. **b** UMAP of myeloid cell populations colored by tissue in integrated scRNAseq data. **c** Barplot of the proportions of myeloid cell populations in synovium and PBMC across the diagnoses. The bar's width corresponds to the number of cells per diagnosis. **d** Bubble plot of myeloid cell clusters marker genes. **e** Dot plot of log2 fold differences in proportions of myeloid cell populations between RA and OA/NIC calculated by permutation test (scProportionTest[44]). **f** Dot plot of log2 fold differences in proportion of myeloid cell populations between PsA and OA/NIC calculated by permutation test (scProportionTest[44]). **g** UMAP of synovial fibroblast cell populations colored by diagnosis in integrated scRNAseq data. **h** Barplot of the proportions of synovial fibroblasts populations across the diagnoses. The bar's width corresponds to the number of cells per diagnosis. **i** Bubble plot of marker genes of synovial fibroblast subpopulations. **j** Dot plot of log2 fold differences in proportions of fibroblasts populations between RA and OA/NIC calculated by permutation test (scProportionTest[44]). **k** Dot plot of log2 fold differences in proportions of fibroblasts populations between RA and PsA calculated by permutation test (scProportionTest[44]).

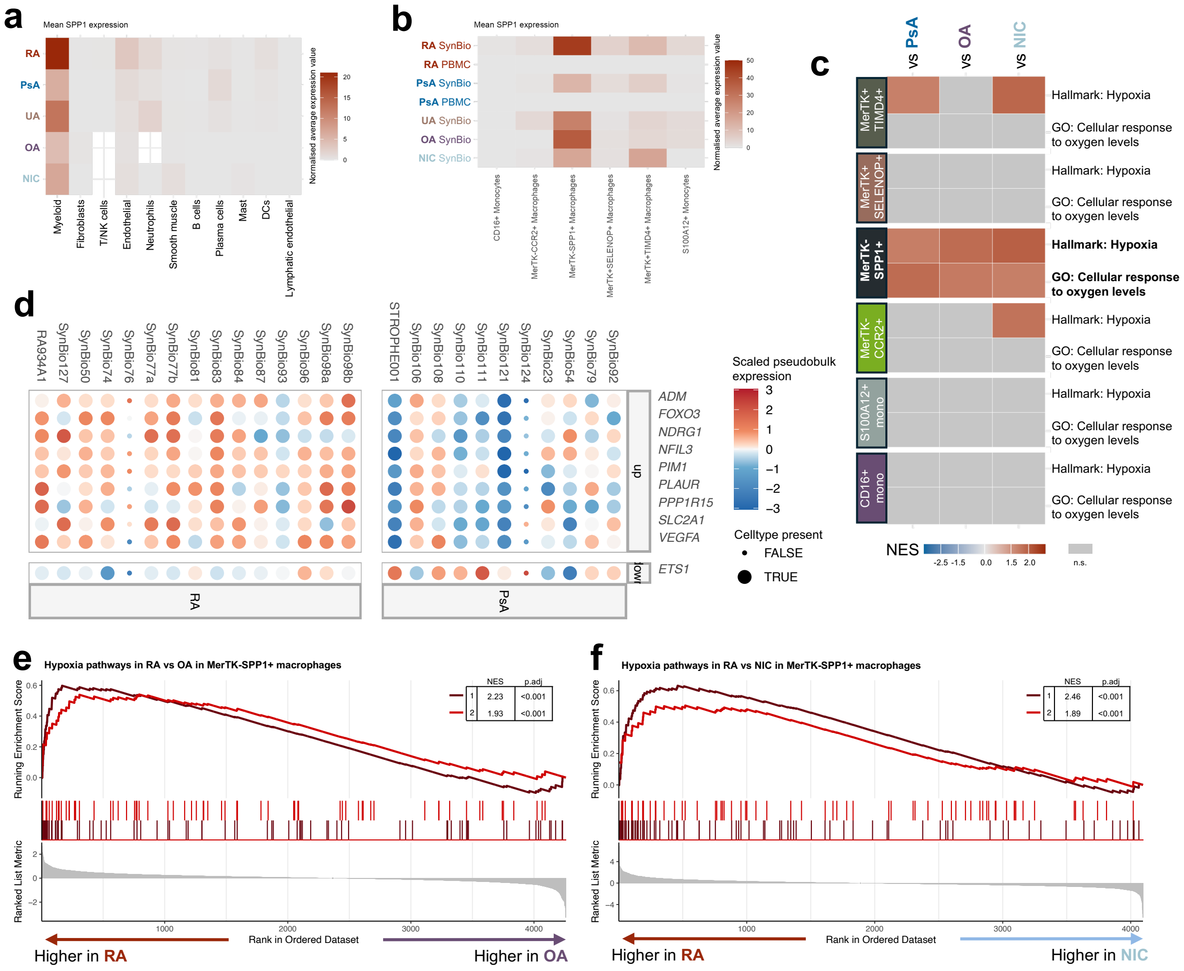

#### Extended Data Figure 3.

**a** Heatmap of *SPP1* expression across major cell types in synovial tissue and blood. **b** Heatmap of *SPP1* expression across myeloid subpopulations in synovial tissue and blood. **c** Heatmap of normalized enrichment score (NES) values of GSEA per myeloid subpopulation in RA versus other diagnosis groups. **d** Dotplot of scaled pseudobulk expression of genes from “Hallmark: Hypoxia” pathway predicted to be up- or down-regulated in MerTK-SPP1$^{+}$ macrophages upon SPP1 interaction with ITGA5 and CD44. **e** GSEA results for pathways up-regulated in MerTK-SPP1$^{+}$ macrophages in RA versus OA (1 – Hallmark: Hypoxia, 2 – GO: Cellular response to oxygen levels). **f** GSEA results for pathways up-regulated in MerTK-SPP1$^{+}$ macrophages in RA versus NIC (1 – Hallmark: Hypoxia, 2 – GO: Cellular response to oxygen levels).

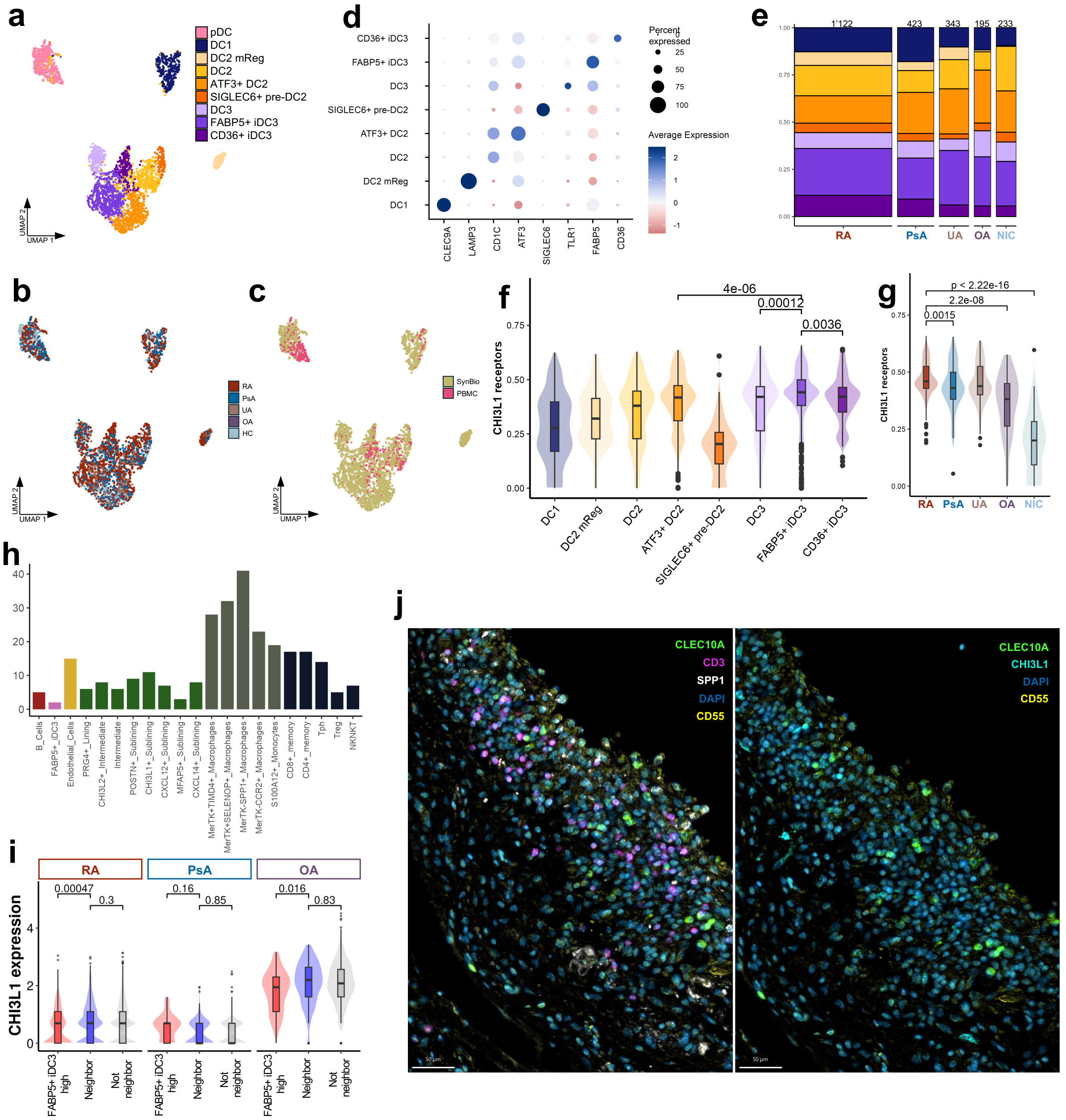

**Extended Data Figure 4.**

**a** UMAP of DC subpopulations in merged scRNAseq data from tissue and peripheral blood. **b** UMAP of DC subpopulations colored by diagnosis in merged scRNAseq data. **c** UMAP of DC subpopulations colored by tissue in merged scRNAseq data. **d** Bubble plot of marker genes of DC subpopulations. **e** Barplot of the proportions of synovial DC populations between the diagnoses. The bar's width corresponds to the number of cells per diagnosis. **f** Boxplot with violin plots of CHI3L1 receptors (*IL13RA2, CD44, TMEM219, LGALS3*) signature values across DC subpopulations (Wilcox test). **g** Boxplot with violin plots of CHI3L1 receptors (*IL13RA2, CD44, TMEM219, LGALS3*) signature values in FABP5$^{+}$ iDC3 across diagnoses (Wilcox test). **h** Number of interaction outgoing from FABP5$^{+}$ iDC3 with the probability > 8% in RA according to CellChat[33]. **i** Boxplot with violin plots of *CHI3L1* expression values across FABP5$^{+}$ iDC3 "neighborhoods" across diagnosis in spatial transcriptomics data. **j** Multiplex IF staining of lining SFs (CD55-positive), DCs (CLEC10A-positive), T cells (CD3-positive), SPP1+ macrophages (SPP1-positive) and CHI3L1$^{+}$ fibroblasts (CHI3L1-positive) in RA synovial tissue.

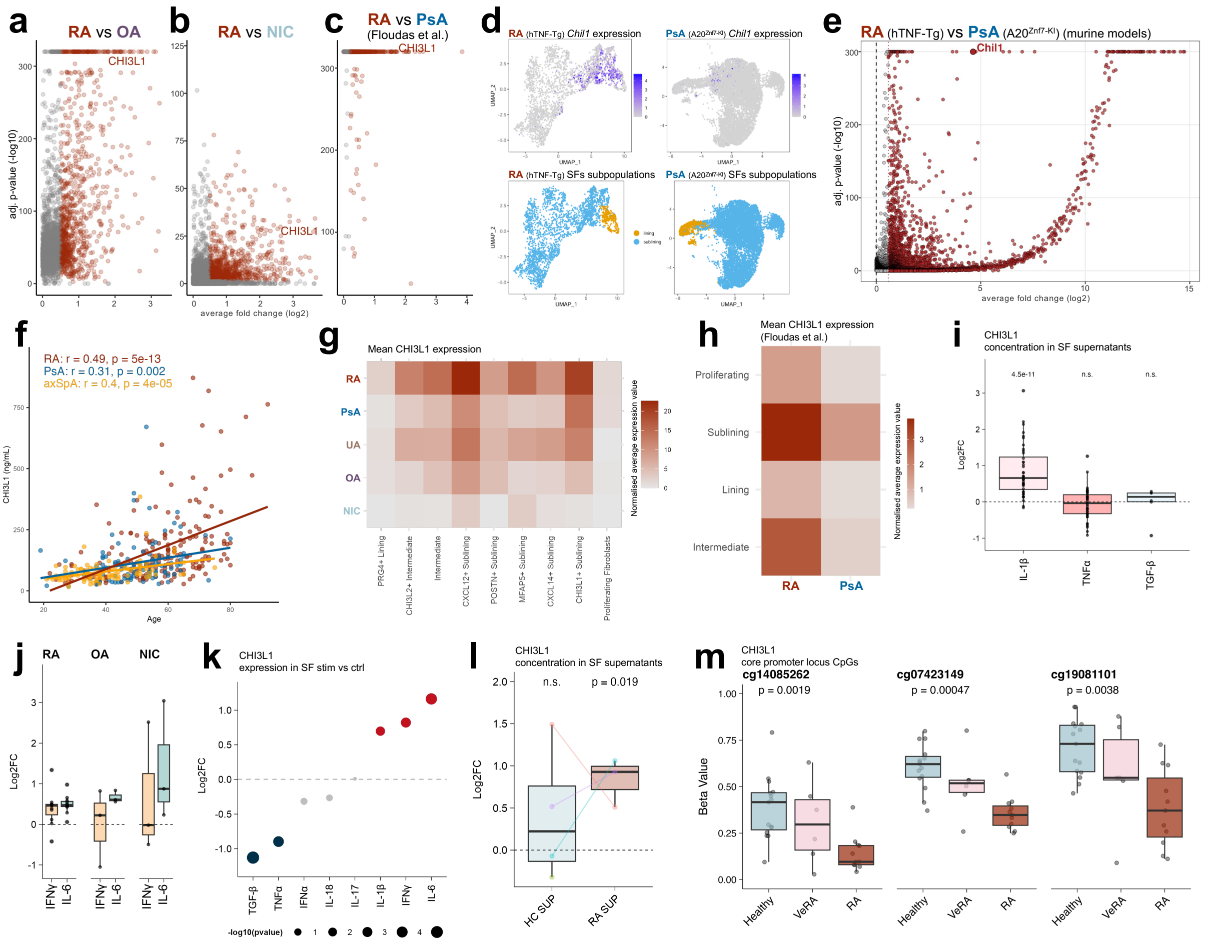


#### Extended Data Figure 5.

**a** Volcano plot of DEGs up-regulated in RA versus OA fibroblasts. **b** Volcano plot of DEGs up-regulated in RA versus NIC fibroblasts. **c** Volcano plot of DEGs up-regulated in RA versus PsA fibroblasts reanalyzed from Floudas *et al.*[11]. **d** Feature plots for *Chil1*expression on the UMAP of SFs derived from a RA-like (hTNF-Tg mouse) and a PsA-like ($A20^{Znf7\text{-}KI}$ mouse) models of arthritis. **e** Volcano plot of DEGs up-regulated in RA-like versus PsA-like SFs. **f** Correlation between CHI3L1 concentration in serum and age by diagnosis (Spearman correlation test) (n = 389: RA = 190, PsA = 99, axSpA = 100). **g** Heatmap of mean *CHI3L1* expression levels across synovial fibroblast subpopulations across diagnoses. **h** Heatmap of mean *CHI3L1* expression levels across synovial fibroblast subpopulations across diagnoses reanalyzed from Floudas *et al.*[11]. **i** Log2 fold change values of CHI3L1 concentration in synovial fibroblast supernatant after stimulation versus control (n = 40) (two-tailed Wilcoxon test). **j** Log2 fold change values of CHI3L1 concentration in synovial fibroblast supernatant after stimulation versus control (n = 16: RA = 11 , OA = 3, NIC = 2). **k** Log2 fold change values of *CHI3L1* expression in synovial fibroblast after different stimulations (re-analyzed from Tsuchiya *et al.*[13]). **l** Log2 fold change values of CHI3L1 concentration in RA synovial fibroblast supernatant after stimulation with supernatants from T-cell obtained from RA patients (RA SUP) and healthy individuals (HC SUP) versus unstimulated (n = 3-4). **m** Beta values of three CpG sites located in the core promoter (cg14085262, cg07423149, and cg19081101) healthy (n = 15), very early RA (VeRA, n = 6), and RA (n = 11) samples (Kruskal–Wallis test).

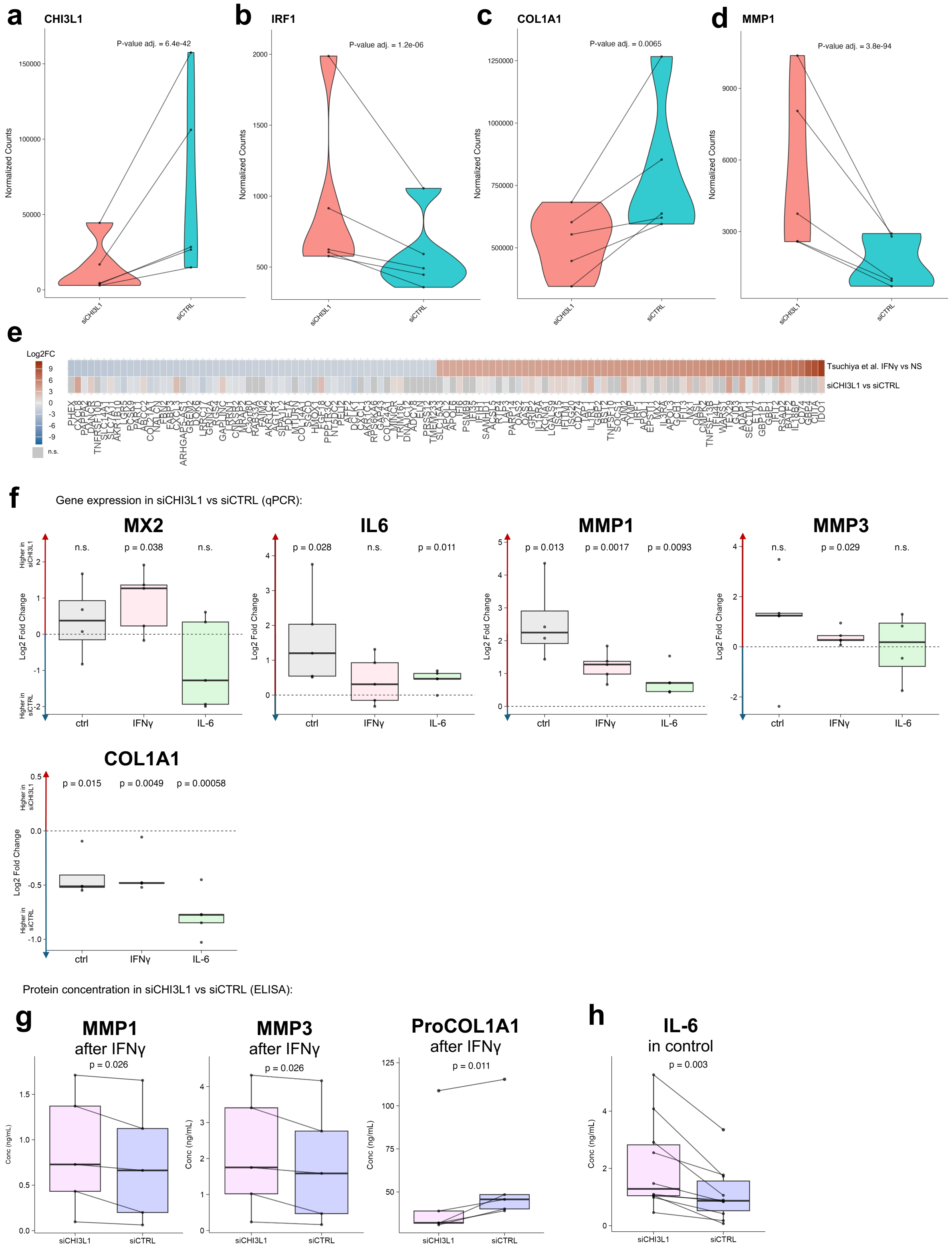

#### Extended Data Figure 6.

**a** *CHI3L1* gene expression in siCHI3L1 and siCTRL (n = 5) in bulk RNA sequencing. **b** *IRF1* gene expression in siCHI3L1 and siCTRL (n = 5) in bulk RNA sequencing. **c** *COL1A1* gene expression in siCHI3L1 and siCTRL (n = 5) in bulk RNA sequencing. **d** *MMP1* gene expression in siCHI3L1 and siCTRL (n = 5) in bulk

RNA sequencing. **e** Heatmap of gene expression log2 fold change in SF stimulated with IFNγ versus non-stimulated (NS) (re-analyzed from Tsuchiya *et al.*[13]) and siCHI3L1 versus siCTRL (n = 5). **f** *MX2*, *IL6, MMP1, MMP3,* and *COL1A1* log2 fold change values of qPCR results in siCHI3L1 (n = 4-5) versus siCTRL (n = 4-5) RA SF (ctrl) and in siCHI3L1 versus siCTRL after stimulations (IL-6 and IFNγ) (one-tailed T-test). **g** MMP1, MMP3, and pro-Collagen I alpha 1 concentration in RA SF supernatants (n = 5) in siCHI3L1 and siCTRL after IFNγ stimulation (one-tailed T-test). **h** IL-6 concentration in RA SF supernatants (n = 10) in siCHI3L1 and siCTRL (one-tailed Wilcox test).

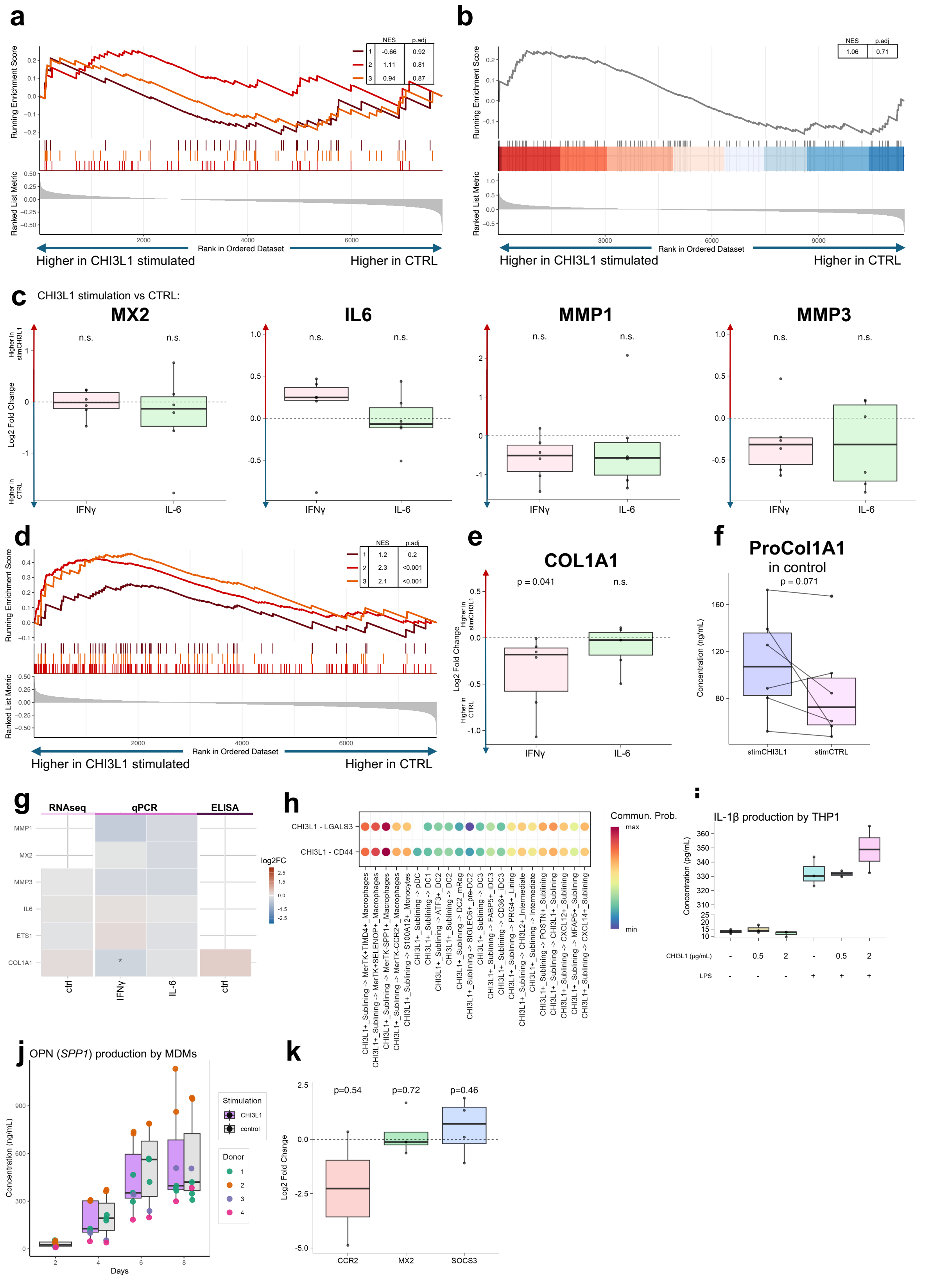


### Extended Data Figure 7.

**a** GSEA results in CHI3L1 stimulated versus non-stimulated (CTRL) SFs for pathways: 1 – Reactome: IFN α/β signaling, 2 – Hallmark: IFNα response, 3 – GO: Response to type I IFN. **b** GSEA result for pathway "Hallmark: Inflammatory response” in CHI3L1 stimulated versus non-stimulated (CTRL) SFs. **c** *MX2*, *IL6, MMP1,* and *MMP3* log2 fold change values of qPCR results in RA SF stimulated with CHI3L1 (n = 5-6) versus non-stimulated (CTRL) (n = 5-6) after stimulations (IL-6 and IFNγ) (one-tailed T-test). **d** GSEA results CHI3L1 stimulated versus non-stimulated (CTRL) SFs for pathways: 1 – Reactome: Collagen formation, 2 – Reactome: Extracellular matrix organization, 3 – GO: Collagen fibril organization. **e** *COL1A1* log2 fold change values of qPCR results in RA SF stimulated with CHI3L1 (n = 5-6) versus non-stimulated (CTRL) (n = 5-6) after stimulations (IL-6 and IFNγ) (one-tailed T-test). **f** Pro-Collagen I alpha 1 concentration in RA SF supernatants stimulated with CHI3L1 (n = 6) and non-stimulated (stimCTRL) (n = 6) (one-tailed T-test). **g** Heatmap of log2 fold change values of gene expression in RA SF stimulated with CHI3L1 versus non-stimulated across different type of analysis (summarized from panels c, e and f). **i** Bubble plot of communication probability via CHI3L1-CD44 and CHI3L1-LGALS3 between CHI3L1$^{+}$ SFs and other SF, myeloid cell and DC subpopulations. **i** IL-1β concentration in THP1 supernatants after stimulation with CHI3L1 and/or LPS (n = 3). **j** OPN (*SPP1*) concentration in MDM supernatants across days after stimulation with CHI3L1 or under control conditions (n donors = 4). **k** *CCR2, MX2, and SOCS3* log2 fold change values of qPCR results in MDMs stimulated with CHI3L1 versus non-stimulated (CTRL) (n = 2-4) (one-tailed T-test).

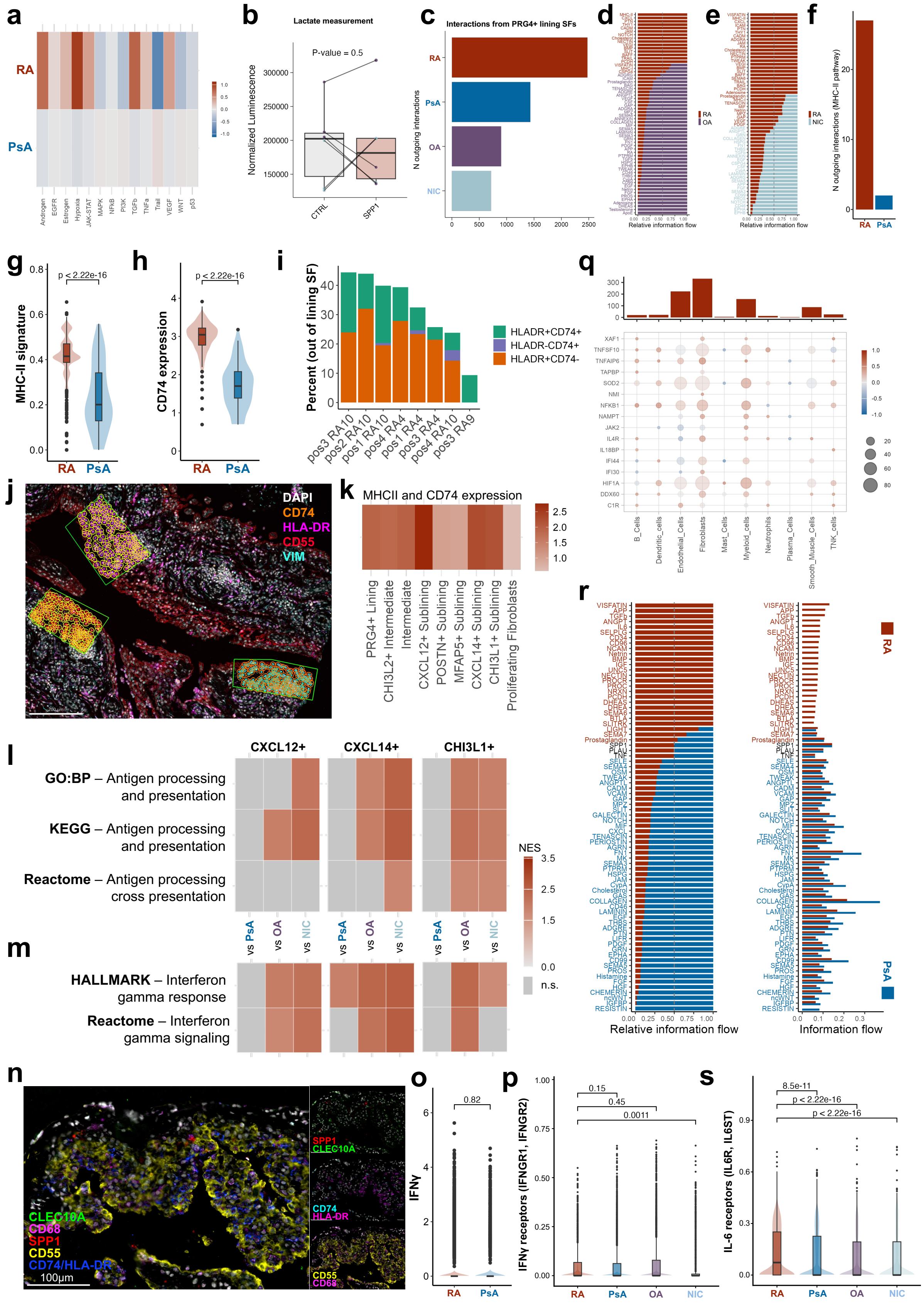

### Extended Data Figure 8.

**a** PROGENy[9] pathway activity scores in lining SFs in RA versus PsA re-analyzed from Floudas *et al.*[11]. **b** Normalized luminescence values reflecting lactate concentration in SF supernatants after SPP1 stimulation and control (n = 6, NIC = 3, OA = 3) (one-tailed paired Wilcoxon test). **c** Number of all interactions outgoing from PRG4$^+$ lining SFs to other cell types across diagnoses. **d** Barplot of cell-cell interactions outgoing from PRG4$^+$ lining SFs in RA compared to OA. **e** Barplot of cell-cell interactions outgoing from PRG4$^+$ lining SFs in RA compared to NIC. **f** Total number of interactions outgoing from PRG4$^+$ lining SFs in RA compared to PsA. **g** Violin plot with bar plot of MHC class II signature (*HLA-DRA, HLA-DQA1, HLA-DQB1, HLA-DQA2, HLA-DQB2, HLA-DPA1, HLA-DPB1* genes) in lining cluster of spatial dataset (two-tailed Wilcox test). **h** Violin plot with bar plot of *CD74* expression in lining cluster of spatial dataset (two-tailed Wilcox test). **i** Immunofluorescence staining analysis of RA synovial tissue showing proportion of HLA-DR-positive and CD74-positive SFs in lining regions. **j** Representative multiplex immunofluorescence staining image and region selection for the analysis in panel *i*. **k** Average expression of HLA class II genes (*HLA-DRA, HLA-DRB5, HLA-DRB1, HLA-DQA1, HLA-DQB1, HLA-DQA2, HLA-DQB2, HLA-DPA1, HLA-DPB1*) and *CD74* across SFs subpopulations of RA patients. **l-m** Heatmap of normalized enrichment score (NES) values of GSEA per SF subpopulation in RA versus other diagnosis groups. **n** Multiplex immunofluorescence staining image of lining SFs (CD55-positive), DCs (CLEC10A-positive), macrophages (CD68-positive), SPP1+ macrophages (double-positive for CD68 and SPP1) and HLA-DR+ fibroblasts (double-positive for HLA-DR and CD74) in RA synovial tissue. **o** *IFNG* expression in all synovial T cells in RA and PsA (Wilcox test). **p** IFNγ receptor signature (IFNGR1, IFNGR2) in PRG4$^+$ lining SFs in RA and PsA (Wilcox test). **q** Upper panel: Number of the interactions significantly correlating (Pearson p-value $< 0.05$) with expression of genes in “Hallmark: IFNγ response” pathway in PRG4$^+$ lining SFs. Lower panel: mean Pearson’s r of correlation between the expression of genes in “Hallmark: IFNγ response” pathway in PRG4$^+$ lining SFs and ligands in corresponding senders and receptors on PRG4$^+$ lining SFs. **r** Barplot of cell-cell interactions incoming to PRG4$^+$ lining SFs in RA compared to PsA. **s** IL-6 receptor signature (IL6R, IL6ST) in PRG4$^+$ lining SFs in RA and PsA (Wilcox test).

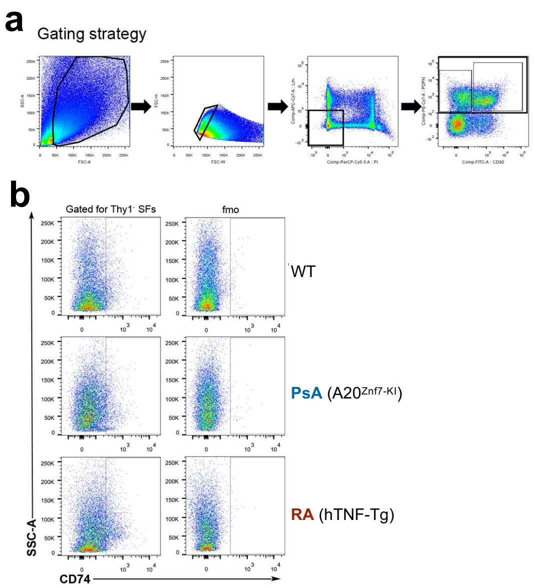


**Extended Data Figure 9.**

**a** Representative flow cytometry gating hierarchy for the identification of CD74⁺ SFs, involving stepwise selection of cells of interest (COI), singlets, and exclusion of CD45⁺ leukocytes, CD31⁺ endothelial cells, and Ter119⁺ erythroid lineage cells prior to gating on CD90⁻PDPN⁺ populations. **b** Representative flow cytometry gating strategy for CD74⁺ lining (CD90⁻) SFs shown for each mouse model. Corresponding FMO controls are displayed for comparison.

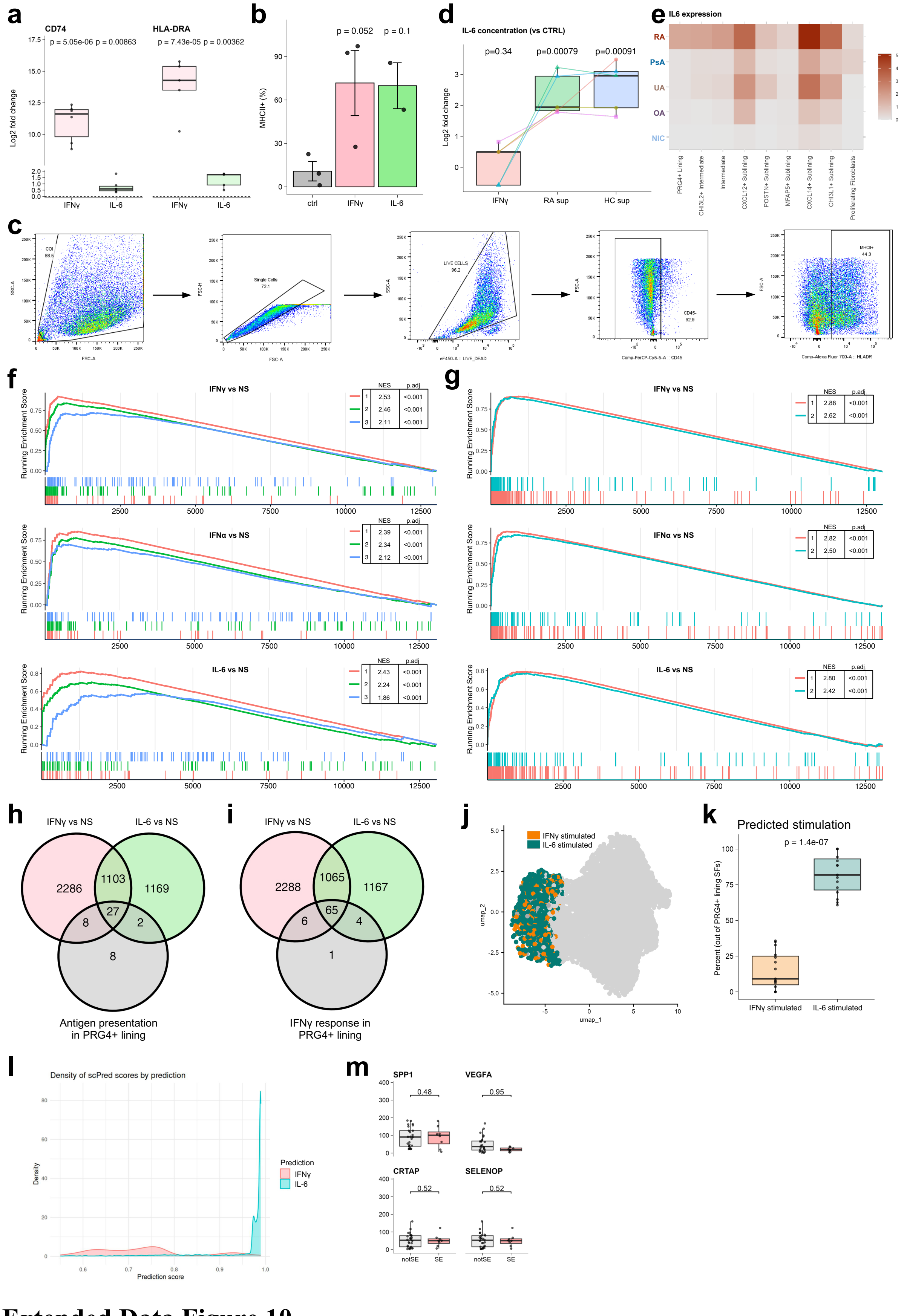


### Extended Data Figure 10.

**a** *CD74* and *HLA-DRA* log2 fold change values of qPCR results in IFNγ and IL-6 stimulated (n = 5-6) versus non-stimulated (n = 5-6) RA SFs (one-tailed T-test). **b** Percentage of MCHII-positive SFs after stimulation with human recombinant IFNγ or IL-6 (one-tailed T-test). **c** Representative flow cytometry gating hierarchy for the identification of $MHCII^+$ SFs, involving stepwise selection of cells of interest (COI), singlets, viable cells, and exclusion of $CD45^+$ leukocytes prior to gating on $MHCII^+$ populations. **d** Log2 fold change of IL-6 concentration in SF supernatant in stimulations with/without blocking versus control (n = 4). **e** Heatmap of mean *IL6* expression levels across SF subpopulations across diagnoses. **f** GSEA results in SFs stimulated with IFNγ, IFNα and IL-6 versus non-stimulated (re-analyzed from Tsuchiya *et al.*[13]) for pathways: 1 – KEGG: Antigen processing and presentation, 2 – GO:BP: Antigen processing and presentation, 3 – Reactome: Antigen processing cross presentation. **g** GSEA results in SFs stimulated with IFNγ, IFNα and IL-6 versus non-stimulated (re-analyzed from Tsuchiya *et al.*[13]) for pathways: 1 – Hallmark: IFNγ response, 2 – Reactome: IFNγ signaling. **h** Venn diagram of genes significantly up-regulated in SFs after IFNγ an IL-6 stimulation (re-analyzed from Tsuchiya *et al.*[13]) and leading edge genes of GSEA analysis of antigen presentation pathways (KEGG: Antigen processing and presentation, GO:BP: Antigen processing and presentation, Reactome: Antigen processing cross presentation) in $PRG4^+$ lining RA versus PsA. **i** Venn diagram of genes significantly up-regulated in SFs after IFNγ an IL-6 stimulation (re-analyzed from Tsuchiya *et al.*[13]) and leading edge genes of GSEA analysis of IFNγ response pathways (Hallmark: IFNγ response, Reactome: IFNγ signaling) in $PRG4^+$ lining RA versus PsA. **j** UMAP of RA SFs with PRG4+ lining colored by predicted stimulations (IFNγ or IL-6). **k** Boxplot of the percentage of $PRG4^+$ lining RA SFs by predicted stimulation. **l** Probability distributions of cells to be classified as IFNγ- or IL-6-stimulated. **m** Boxplot with the numbers of peptides predicted to bind to HLA class II alleles. SE – shared epitope alleles (Wilcox test).
